# Incorporating climatic extremes using the GEV distribution improves SDM range edge performance

**DOI:** 10.1101/2024.10.14.618163

**Authors:** Ward Fonteyn, Josep M. Serra-Diaz, Bart Muys, Koenraad Van Meerbeek

**Author notes:** **Author Note** Ward Fonteyn Conceptualization-Equal, Formal analysis-Lead, Methodology-Lead, Visualization-Lead, Writing – original draft-Lead, Writing – review editing-Equal Josep M. Serra-Diaz Conceptualization-Equal, Methodology-Supporting, Supervision-Equal, Writing – review editing-Equal Bart Muys Conceptualization-Supporting, Supervision-Supporting, Writing – review editing-Supporting Koenraad Van Meerbeek Conceptualization-Equal, Methodology-Supporting, Supervision-Equal, Writing – review editing-Equal. Code and data are available at https://github.com/wardfont/JoBG-sdm-extremes.

## Abstract

**Aim:** The changing frequency and intensity of climatic extremes due to climate change can have sudden and adverse impacts on the distribution of species. While species distribution modelling is a vital tool in ecological applications, current approaches fail to fully capture the distribution of climatic extremes, particularly of rare events with the most disruptive potential. Especially at the edges of species’ ranges, where conditions are already less favourable, predictions might be inaccurate when these extremes are not well represented.

**Location:** Europe

**Taxon:** Tree species

**Methods:** We present a novel approach to integrate extreme events into species distribution models based on the generalised extreme value (GEV) distribution. This distribution, following from the extreme value theory has been established as a valuable tool in analysing climatic extremes, both in an ecological context and beyond. The approach relying on the GEV distribution is broadly applicable, readily transferable across species and relies on widely available data. We demonstrate the efficacy of our approach for 28 European tree species, illustrating its superior ability to fully capture the distribution of climatic extremes compared to state-of-the-art methods.

**Results:** We found that incorporating parameters on climatic extremes derived from the GEV distribution increased model performance (AIC_model_) and characterized range edges more accurately (AUC_edge_) compared to competing approaches. However, general AUC values were only marginally increased across the species and study period analysed. Overall, the GEV model predicted a narrower niche for the species included in this study.

**Main conclusions:** Incorporating climatic extremes can impact spatial predictions of species distribution models, especially at range margins. We found that using the GEV distribution to characterise extreme variables in SDMs yields the best performance at these distribution edges. Given the importance of range edges for species conservation, a detailed inclusion of extremes in SDMs employed for those applications will help ensure robust conclusions.

## Introduction

The study of species distributions plays a crucial role in ecological applications, including testing ecological theories, assessing the impact of climate change, and developing conservation and risk management policies (Srivastava et al., 2019). The prediction and understanding of species distributions often rely on species distribution models (SDMs), which link presence and absence records of a species with the environmental conditions at those locations (Elith & Leathwick, 2009; Franklin, 2010; Guisan et al., 2017). These models not only aid in understanding and predicting present-day distributions, but are also used to forecast range shifts under future climate scenarios. Accurate modelling of species distributions is essential for these applications. We argue in this paper that climatic extremes hitherto are not fully captured in SDMs, even when using state-of-the-art approaches.

In what follows, we define climatic extremes based on the distribution of a climatic variable, as well-defined, but historically rare events. It is widely accepted that climatic extremes affect species and their distribution (Seabrook et al., 2014; Lynch et al., 2014; Germain & Lutz, 2020). Climatic extremes impact organisms non-linearly, exhibiting threshold behaviour (Mitchell et al., 2014). An example is xylem cavitation in vascular plants, which causes abrupt loss of conductivity beyond a certain level of drought stress (McDowell et al., 2008; Choat et al., 2012; Feng et al., 2021). Not only an increase in intensity, but also in frequency of extremes can enlarge the impact on a system (Dreesen et al., 2014; Sánchez-Pinillos et al., 2022). Both can individually or jointly inhibit regeneration or increase mortality, potentially leading to local extinction. In some conifer species for example, previous cavitation can lead to an increased vulnerability to future cavitation (Feng et al., 2021). Identifying a single fixed threshold for a variable representing an extreme, that captures the response of a species to that extreme and which holds for a species across individuals and locations is therefore unlikely. Instead, the full probability density distribution of variables representing climatic extremes should be considered, capturing all combinations of intensity and frequency which may impact a species. Due to the process by which it arises, the distribution of those variables representing extremes is not expected to follow a Gaussian distribution, but rather a generalised extreme value (GEV) distribution. According to the extreme value theory, the GEV distribution results from taking the maximum of a finite number of independent and identically distributed random variables (Katz et al., 2005). Unlike the Gaussian distribution, this GEV distribution is asymmetric and characterised by a skew.

Given the importance of climatic extremes and their potential to influence species’ distribution, a need exists for a method which effectively incorporates their characteristics in SDMs. Some studies incorporate climatic extremes in SDMs by utilising highly detailed data such as daily environmental time series, at times coupled with a method tailored to a specific species, e.g. selecting predictor variables based on expert knowledge of a species’ reproduction or survival (Reside et al., 2010; Morán-Ordóñez et al., 2018; Feldmeier et al., 2018). Other studies identify thresholds for certain variables representing climatic extremes and use them to model and understand the effects of extremes on species distributions (Seabrook et al., 2014; Cavanaugh et al., 2015). Although these approaches can benefit SDMs, the data they rely on is difficult to attain. That is, a detailed understanding of the effects of extremes is often lacking for many species, and it can only be obtained experimentally with considerable effort. Additionally, the use of context-dependent and species-specific knowledge limits the transferability of these methods across species, study areas and climate conditions. Instead, we strive towards a broadly applicable method in this paper, which utilises widely available data and information, and which does not suffer from the same limitations regarding transferability. In the following paragraphs, we identify three existing approaches for SDMs which fulfil these criteria and discuss their shortcomings in capturing the full probability density distribution of climatic extremes.

Variables representing climatic extremes are already part of the bioclimatic variables provided by datasets like WorldClim (Fick & Hijmans, 2017) and therefore many studies include them in SDMs (Booth et al., 2014; Bradie & Leung, 2017). Yet, it is often only the long-term mean of climatic extremes, i.e. the mean extreme intensity, that is used, even in studies explicitly considering extremes (e.g. Bateman et al., 2012; Seabrook et al., 2014; Feldmeier et al., 2016; Morán-Ordóñez et al., 2018; Feldmeier et al., 2018).

Although these means may serve as proxies for physiologically relevant phenomena such as absolute extremes (Körner, 2021), they are severely lacking discriminating power when considering the skewed distribution extremes exhibit. As this ubiquitous approach relies only on the mean, it is henceforth referred to as the Mean approach (Fig. 2).

Zimmermann et al. (2009) included interannual variability of climatic variables, expressed as standard deviation, in addition to their mean to better account for climatic extremes in SDMs. This approach resulted in models with improved predictive power and corrected spatial predictions. They argued that extremes are better represented through variability as opposed to extreme event indices based on thresholds or return periods, as these indices are highly correlated with mean and variability, species-specific, and subject to change. However, characterising extremes solely through interannual variability presents challenges, especially under changing conditions. Although changing variability may alter extremes, it is not the only cause, and both concepts should be considered separately (Bailey & van de Pol, 2016). A change in variability can change the likelihood of extreme events with a certain intensity, but so can a shift in mean or a change in skew while variability remains unchanged. This approach, which relies only on mean and variability, is not adept at capturing changes in skew. As it is equivalent to characterising a normal distribution, it is therefore referred to as the Normal approach (Fig. 2).

Stewart et al. (2021) introduced a third approach using the intensity for specific return periods, derived from empirical quantiles of monthly climatic extreme variables, in addition to their mean to account for climatic extremes in SDMs. The intensity for a specific return period, say 15 years, is the extreme intensity expected to occur or be exceeded every 15 years. They constructed a separate model for each of several return periods, retaining the best-performing model. Their analysis also improved the predictive power of models and resulted in differing spatial predictions. Using empirical quantiles has some limitations, however. Since empirical quantiles are an interpolative method, the highest extreme intensity that can be estimated, is limited to the maximum intensity observed in the dataset. As only a short period is typically considered when characterising climate, this limits the ability to capture rare high-intensity extremes. Additionally, as the empirical density function is not based on a theoretical density function, any quantile estimate is either discontinuous or based on linear interpolation between observations corresponding to adjacent empirical quantiles (See *quantile* function in the *stats* R package (R Core Team, 2022)). This derivation from a limited sample severely biases the estimated extreme intensity for a certain return period. Finally, similarly to the discussion on thresholds, it is unlikely that one single return period will accurately capture the effect of climatic extremes on a species across individuals, space, and time. As such, the selected return period yielding the best model for the present does not necessarily result in the best model for future climate scenarios. This approach is referred to as the Quantile approach (Fig. 2).

When applied to variables representing extremes, these three approaches fail to capture the skew of these variables in their models, which have only one or two parameters per variable. We cannot ignore this skew and assume it to be constant, or assume the differences in skew to be negligible across space and time. Climate change, for example, can affect the skew, distorting the current relationships between average conditions and extremes (Zwiers et al., 2013; Seneviratne et al., 2021).

In this paper, we propose a method for incorporating extremes in SDMs that is transferable and broadly applicable. Our approach is based on the GEV distribution (the GEV approach), and accounts for the skewed nature of variables representing climatic extremes (Fig. 2). The GEV distribution is often employed when analysing extremes, i.e. temperature extremes (Wehner et al., 2020) and river discharge extremes (Kousar et al., 2020). It has also proved to be a valuable tool when analysing the impact of extremes in an ecological context (D. C. Rypkema et al., 2019). Like a normal distribution is characterised by the mean and standard deviation, the GEV distribution is characterised by the location (*µ*), scale (*σ*) and shape (*ξ*) parameters, determining the centre, spread and shape of the distribution, respectively. Three types of GEV distribution are distinguished based on the shape parameter (Fig. 1): the bounded Weibull distribution (*ξ* < 0), the light-tailed Gumbel distribution (*ξ* = 0) and the heavy-tailed Fréchet distribution (*ξ* > 0). If the minimum rather than the maximum is of interest, values are multiplied by -1 and results for the parameters are backtransformed. Many variables describing climatic extremes arise from a so-called block-maxima approach, by taking the maximum across a set amount of time, for instance one year. As a result, those variables are expected to exhibit a GEV distribution rather than a Gaussian distribution. In what follows, we refer to such variables as *extreme variables*. A more in-depth overview of the extreme value theory in ecological applications is given by Katz et al. (2005) and more recently by (D. Rypkema & Tuljapurkar, 2021).

**Figure 1.**
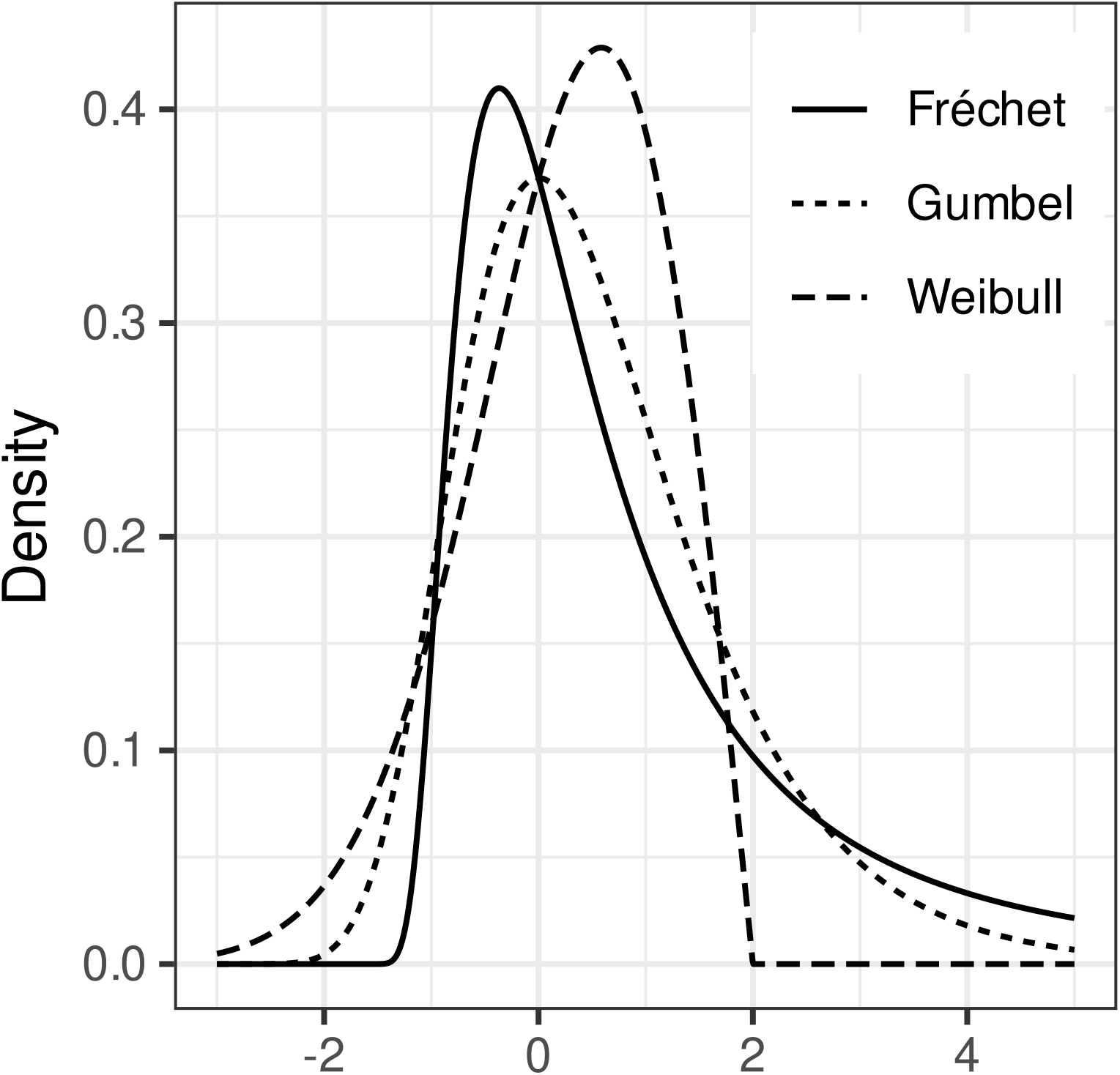
GEV distribution forms arising from µ = 0, σ = 1 and different values for the shape parameter: ξ = -0.5 (Weibull distribution), ξ = 0 (Gumbel distribution), and ξ = 0.5 (Fréchet distribution).

**Figure 2.**
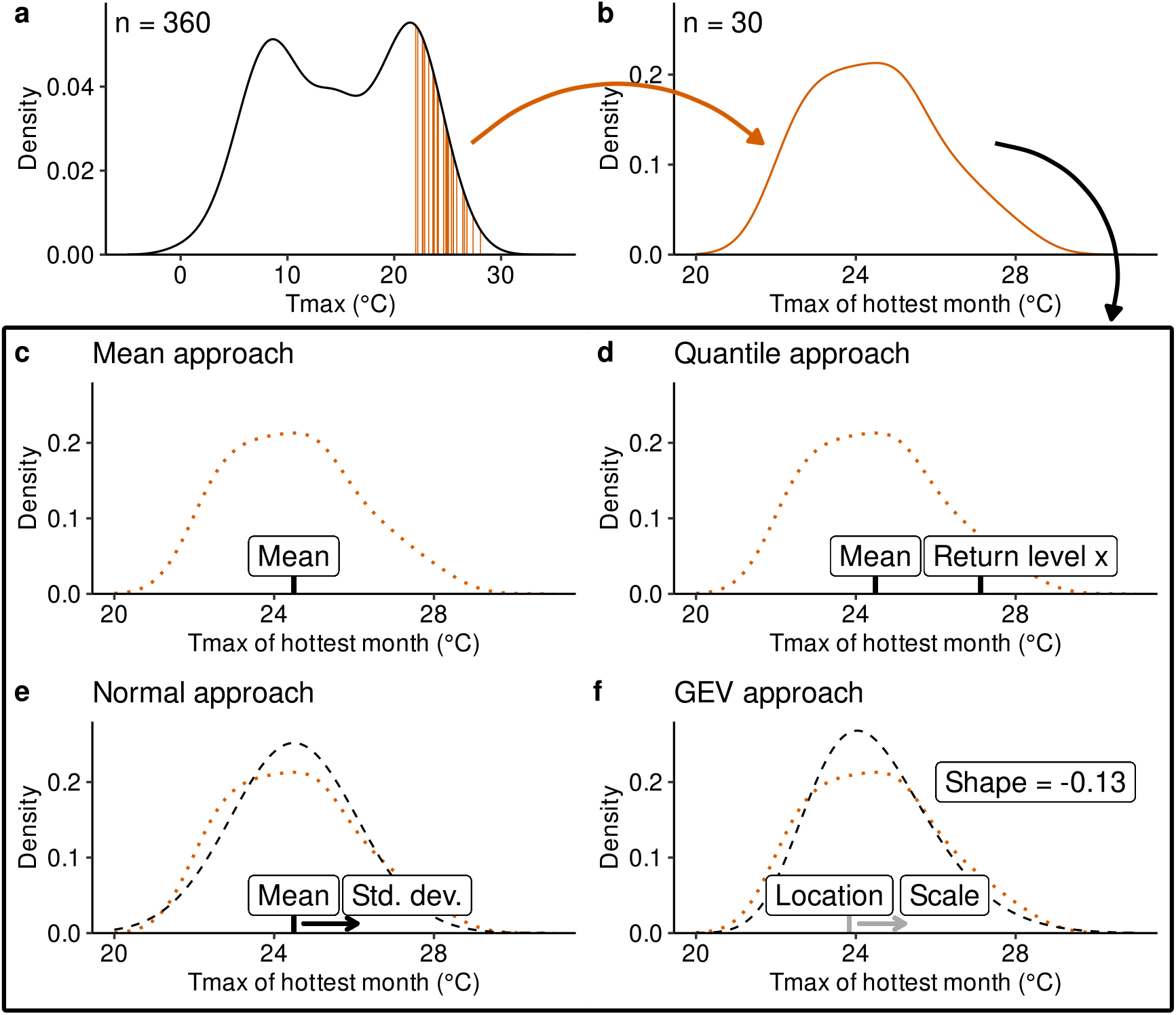
Graphical overview of the discussed approaches to characterise the distribution of variables representing climatic extremes. Here, the climatic extreme of interest is the maximum temperature of the hottest month. The probability density distribution of the monthly maximum temperature is given (a) with the maximum temperatures of the warmest month for each of the 30 years indicated in vertical lines (orange), from which the probability density distribution of the maximum temperature of the hottest month (TxHm) is derived (b). The derived parameters used in different approaches are shown, together with the theoretical distribution they may characterise as a dashed line, and the probability density curve of TxHm as a dotted line (c-f). Unlike the GEV approach, the Mean, Quantile and Normal approaches are unable to capture the skewed nature of the climatic extreme variable’s distribution using only one or two parameters. Data is derived from the TerraClimate dataset for the pixel at 4.70° E, 50.88° N.

In the context of yearly extremes, *µ* is the intensity of an extreme with a return period of *e* years, with *e* being Euler’s number or ∼2.718. In other words, *µ* is the ∼36.79^th^ percentile. It only corresponds with the mode when *ξ* is zero. *σ* characterises the spread of extreme intensities, with higher *σ* indicating wider spread. Finally, *ξ* denotes the likelihood of very intense extremes. When *ξ* is negative, the right tail of the GEV distribution is bounded, meaning that very intense extremes after a certain cut-off are not expected to occur, with more negative values for *ξ* corresponding to a lower, less intense cut-off. When *ξ* is equal to zero or positive, the right tail is not bounded, and the probability of very intense extremes increases with the value of *ξ*. Although the extreme value theory assumes independent and identically distributed random variables, it has been shown to hold even for stationary random variables where independence is not fulfilled (Coles, 2001). Additionally, where stationarity is not met, the GEV parameters can be set to vary over time, e.g. a linear trend in the location parameter while fitting the GEV distribution results in different estimates for the scale and shape parameters (Coles, 2001).

The GEV approach presented in this paper relies on widely available data, without additional species-specific knowledge. In a case study based on 28 European tree species with diverging ecological traits, we explore the ability of predictor variables derived from the GEV distribution of several climatic extreme variables to improve species distribution modelling beyond the other approaches discussed above.

## Methods

### Case study: modelling European tree species using three climatic extreme variables

In this analysis, our aim is to assess whether including additional details of climatic extremes improves the predictive performance of SDMs, and which approach yields most benefit. We therefore compare models constructed following the Mean approach to those following the Quantile, Normal and GEV approaches. We model the distribution of 28 European tree species with differing ecological traits. The different approaches are applied to three extreme variables that influence a species’ distribution through acute stress: heat, cold and precipitation deficiency (L. D. L. Anderegg & Lambers, 2016; Ruehr et al., 2019; Körner, 2021). In addition to the parameters for these three climatic extreme variables, bioclimatic variables are included in the SDMs to adhere to common practices in variable selection. Besides the predictor variables specific to each approach, all modelling choices are kept identical across approaches to provide a standardised basis for comparison. Models for all approaches are based on the same data, with the only difference being the detail with which the three extreme variables are characterised. The study area was defined as Europe up to a longitude of 34° East, the British Isles, and Mediterranean islands excluding Malta and Cyprus. Although this example was conducted on tree species, the employed methods are transferable to all analyses where extreme variables are of interest.

### Climate variables

Monthly time series of maximum temperature, minimum temperature and precipitation were acquired from the TerraClimate dataset (Abatzoglou et al., 2018). This dataset combines WorldClim climatological normals with monthly time series resulting in a 1/24degree spatial resolution (∼3 km at a 50° latitude) at a monthly frequency, available from 1958 to 2021 at the time of writing. The climate for the 30-year period from 1986 to 2015 was considered for this analysis, corresponding to the reference period in the TerraClimate dataset. We chose a 30-year period to align with standard practices (Fig. 3).

**Figure 3.**
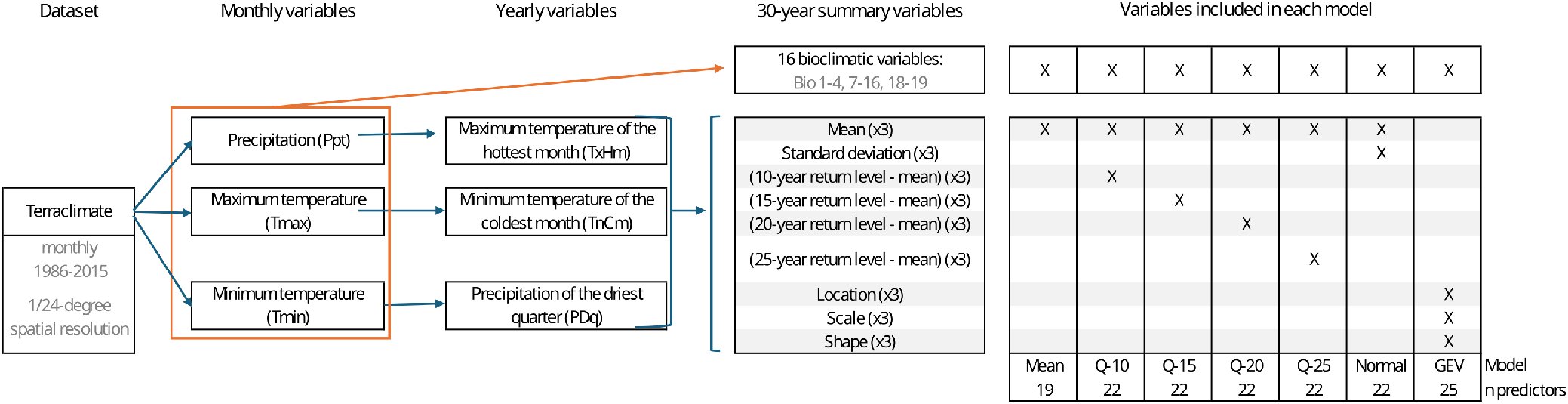
Schematic overview of the climate data used in the analysis, the climate variables that were derived from them, and in which model approaches the derived variables were included. The “x3” signifies the parameters for each of the TxHm, TnCm and PDq.

From these monthly climate data, the three extreme variables representing heat, cold and precipitation deficiency were derived: maximum temperature of the hottest month (*TxHm*), minimum temperature of the coldest month (*TnCm*) and precipitation of the driest quarter (*PDq*). This was achieved by using the block-maxima approach with yearly blocks of 12 months, i.e. taking yearly maxima for the maximum temperature, and yearly minima for both the minimum temperature and the 3-month rolling window precipitation sum. For minimum temperature, yearly samples were centred around the winter (starting from July and ending in June), resulting in 29 yearly minima. For precipitation, the first year was dropped as no rolling sum could be calculated for the first two months, also resulting in 29 yearly minima.

The different parameters needed by the modelling approaches were now calculated for each of the TxHm, TnCm and PDq. For each pixel, the mean, standard deviation, and empirical quantiles corresponding to a return period of 10, 15, 20 and 25 years expressed as anomalies to the mean, were calculated. Additionally, the GEV parameters for each extreme variable were estimated by the maximum likelihood method using the *fevd* function of the *extRemes* R package (Gilleland & Katz, 2016). As these parameters characterise a 30-year period, we assume the climate in this period to be stationary, in which case the extreme value theory holds as mentioned above. Although this assumption is here explicitly stated for the calculation of the GEV, the same assumption is implicitly made when summarising the 30-year period using the Mean, Quantile and Normal approaches.

Only the Normal and GEV approaches specify a distribution for the extreme variables using the derived parameters. To assess which distribution had a superior fit for the three extreme variables, the Akaike information criterion of the distributions fitted to TxHm, TnCm and PDq values for each pixel was calculated (AIC_distr._). A lower AIC_distr._ indicates an improved fit, translating into a better characterisation of the extreme variables’ distribution while penalising the increased complexity of the GEV distribution.

To construct the SDMs, we additionally used the 19 bioclimatic variables as defined in Fick & Hijmans (2017). The description of all 19 bioclimatic variables is given in Appendix A. Note that the mean of the *TxHm, TnCm* and *PDq*, as calculated above, correspond to bioclimatic variables 5, 6 and 17, respectively. The remaining 16 bioclimatic variables were calculated based on the monthly maximum temperature, minimum temperature and precipitation using the *biovars* function of the *dismo* package (Hijmans et al., 2022).

### Species records

The EU-Forest dataset provides data on forest plots in Europe based on systematic sampling in national forest inventories and research networks, aggregated to a 1 km grid (Mauri et al., 2017). Being based on national forest inventories, this data includes observations for trees of all age classes, including seedlings. The species records cover the climatic gradient of Europe well. Based on the availability of ample records (> 4000 presences), 28 tree species were considered. As only common tree species were included, we assume that they would be identified if present in the systematic sampling. Therefore, any site where these species are not noted as present, was taken as an absence. For each species, only absences within 500 km of a presence were retained to exclude false absences from areas not historically accessible to the species. The remaining presence and absence records were spatially thinned to the 1/24-degree resolution of the climate data to reduce the spatial bias.

### Species distribution model fitting

Per species, the presence-absence dataset was first split into ten random folds. For each of these folds, a separate model was constructed. This random assignment reduces the number of observations per model, decreasing computation time, and introduces stochasticity between the random folds, aiding in the interpretation of the results for model performance. Subsequently, spatially blocked partitioning based on a square grid, which minimised spatial autocorrelation of all predictor variables across species records, was performed on each of those random folds using the *part_sblock* function from the *flexsdm* R package (Velazco et al., 2022). The block size with minimal spatial autocorrelation was selected from 30 block sizes ranging from 25 to 300 times the climate data spatial resolution. The spatial autocorrelation was calculated based on a randomly assigned 10% of species records. Starting at ten spatially blocked partitions, the number of partitions was reduced if needed until each partition contained at least 1% of total presences. These spatially blocked partitions were used for cross validation of the models.

Seven models spanning the four approaches were constructed. The Mean model incorporated only the means of the extreme variables. Four Quantile models included both the mean and the anomaly to the mean of the expected extreme intensity for either the 10-, 15-, 20- or 25-year return period. The Normal model included the mean and standard deviation. Finally, the GEV model included the location, scale, and shape of the GEV distribution fitted to each extreme variable. In addition, all models included the 16 bioclimatic variables.

Species distributions were modelled using generalised additive models (GAMs) with a binomial distribution. GAMs are a performant algorithm (Valavi et al., 2022), which allows for the calculation of the AIC of the fitted model (AIC_model_). The AIC_model_ gives essential information on model fit when comparing models with a different number of predictor variables. The GAMs included a linear combination of smooth terms for all predictors and were constructed using *gam* function of the *mgcv* R package (Wood, 2011). The default methods of this function were maintained, with smooth terms as penalised regression splines with 10 basis functions and smoothing parameter estimation using the generalised cross validation criterion. Model terms were not penalised. We document the models from this analysis using the ODMAP protocol (Zurell et al., 2020) in the Appendix G.

### Model evaluation

As Fourcade et al. (2018) show, the evaluation and comparison of species distribution models is a complex topic which should best be supported by multiple metrics. Therefore, model fit was evaluated using the AIC_model_, which allows us to compare fit across models with differing numbers of predictors. It acts as an estimate of the out-of-sample deviance and therefore of the model predictive performance (McElreath, 2016). As AIC_model_ can only be used for relative assessments, model predictive performance was further assessed using the spatially blocked design with the threshold-independent area under the receiver operating characteristic curve (AUC).

Extremes are expected to exert influence mostly at the species’ distribution edges (Zimmermann et al., 2009). We identify these distribution edges using the Mean model predictions for each species and random fold separately. Here, edge pixels were taken as those pixels with a predicted probability of occurrence by the Mean model lower than the threshold which maximises the sum of sensitivity and specificity, i.e. the true skill statistic (TSS) maximising threshold, and higher than 0.025. The lower limit was set to ensure the edge region was limited. We then calculate the AUC_edge_ for each model based only on those species presences and absences which are located in the edge region. This AUC_edge_ gives us a measure for how well each model performs within the distribution edges.

Differences in model evaluation metrics between the constructed SDM approaches were assessed using generalised linear mixed-effects models using the *glmmTMB* function from the identically named R-package (Brooks et al., 2017). Mixed-effects models were chosen because they can account for the nested structure of the data (i.e. performance metric values nested within random folds and random folds within species). Three models were constructed, including the modelling approach as a categorical predictor variable and each performance metric separately as response variable, while the species and random folds were included as nested random variables. For the AIC_model_, an unbounded metric, a standard linear mixed-effects model with Gaussian distribution was used. For AUC and AUC_edge_, which are bounded between 0 and 1, generalised linear mixed-effects models with a beta distribution and a logit link function were used instead to account for this bounded nature. Each model included 1960 observations across 280 random folds nested within 28 species. From these mixed-effects models, contrasts in estimated marginal mean model performance between the employed SDM approaches were assessed using the *emmeans* R-package (Lenth, 2022) with a Benjamini-Hochberg adjustment.

### Spatial predictions

To analyse differences in continuous spatial predictions made by different modelling approaches, the models for the ten random folds were combined into one unweighted mean ensemble model for each species using the *fit_ensemble* function of the *flexsdm* R package. To avoid interpreting extrapolative error, only spatial predictions within the species’ 500 km calibration area were evaluated. The spatial predictions for the GEV ensemble model were contrasted against those for the Mean ensemble model. This contrast was quantified in two ways. The absolute difference between predictions highlights those locations where the choice of modelling approach has a large impact. However, only locations where large absolute differences are possible can experience such large impacts. If the Mean approach predicts a low probability of occurrence (e.g. 0.1), the prediction by the GEV approach can only decrease by that much (−0.1), while much larger increases are possible (up to 0.9). Vice versa, for high predictions by the Mean approach (e.g. 0.9), only a limited increase is possible (0.1) while a much larger decrease can take place (up to -0.9). The relative difference between the predictions of both approaches therefore highlights the areas where the increase or decrease is large compared to the maximum possible increase or decrease. To calculate the relative differences between the GEV approach and the Mean approach, decreases in probability of occurrence for the GEV approach compared to the Mean approach are divided by the predicted probability of occurrence for the Mean approach (predicted_Mean_), while increases are divided by (1-predicted_Mean_). All spatial predictions were based on the same 1/24-degree climate raster data as used to train the SDMs.

## Results

### Species-independent fit of theoretical distribution to the extreme variables

The comparison of the AIC_distr._ between the normal distribution and the GEV distribution reveals the superior ability of the GEV distribution to characterise the three extreme variables in the European study area (Fig. 4). The GEV distribution yields a lower AIC_distr._ and therefore an improved fit over the normal distribution in most of the European study area for all three extreme variables. Only for the two extreme variables related to temperature does the GEV distribution not surpass the normal distribution in some areas, most notably the Mediterranean region in the case of TxHm. For the PDq, the GEV distribution is preferred across the board.

**Figure 4.**
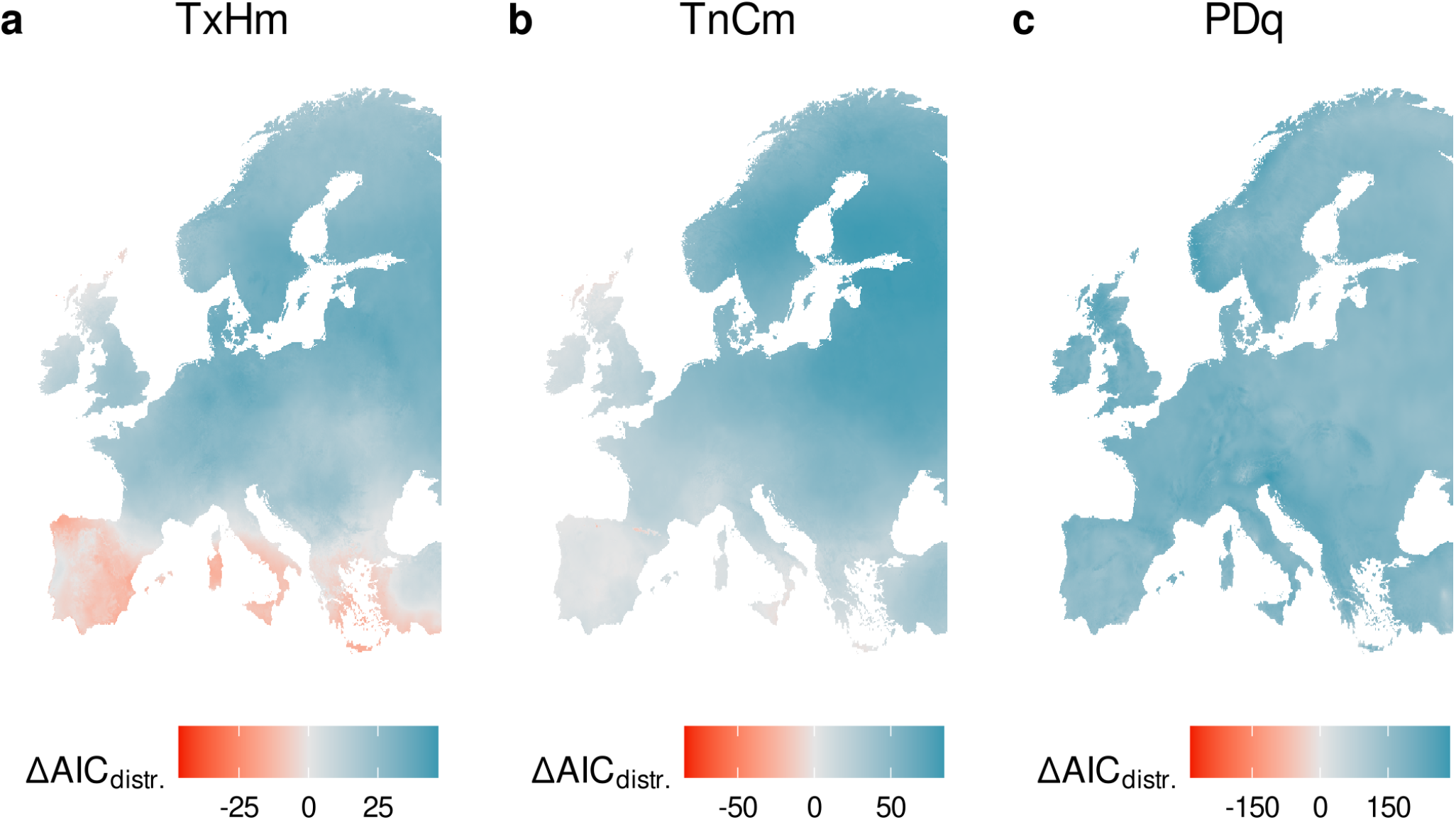
The GEV distribution provides a better fit in most of the European study area compared to the normal distribution for the three extreme variables. The difference in AIC_distr._ between the normal distribution and GEV distribution fit (AIC_distr._(N) − AIC_distr._(G)) is shown for maximum temperature of the hottest month (TxHm), minimum temperature of the coldest month (TnCm) and precipitation of the driest quarter (PDq). Positive values indicate a lower AIC_distr._ for the GEV distribution, corresponding to a better fit while penalising for the increased complexity of an additional parameter. These calculation were performed on 1/24-degree spatial resolution climate data rasters.

### Model predictive performance

Compared to the Mean model containing only the bioclimatic variables and the mean of the extreme variables, all other models including additional characteristics for those extreme variables showed a lower AIC_model_, with the GEV model showing a significantly lower AIC_model_ compared to all other models (Fig. 5). Average species cross-validated AUC of the Mean model ranged from 0.63 to 0.96, with only two species with a value lower than 0.75. The GEV model, the Normal model and two of the Quantile models still showed a significant improvement in AUC. However, no significant differences existed between those better-performing models. Compared to the small effect sizes seen for the overall AUC, the AUC_edge_ shows much larger effect sizes. From the AUC_edge_, for which the average for the Mean model ranged from 0.52 to 0.82, the same conclusions as for AIC_model_ can be drawn, with all models outperforming the Mean model and the GEV model performing significantly better than all other models.

**Figure 5.**
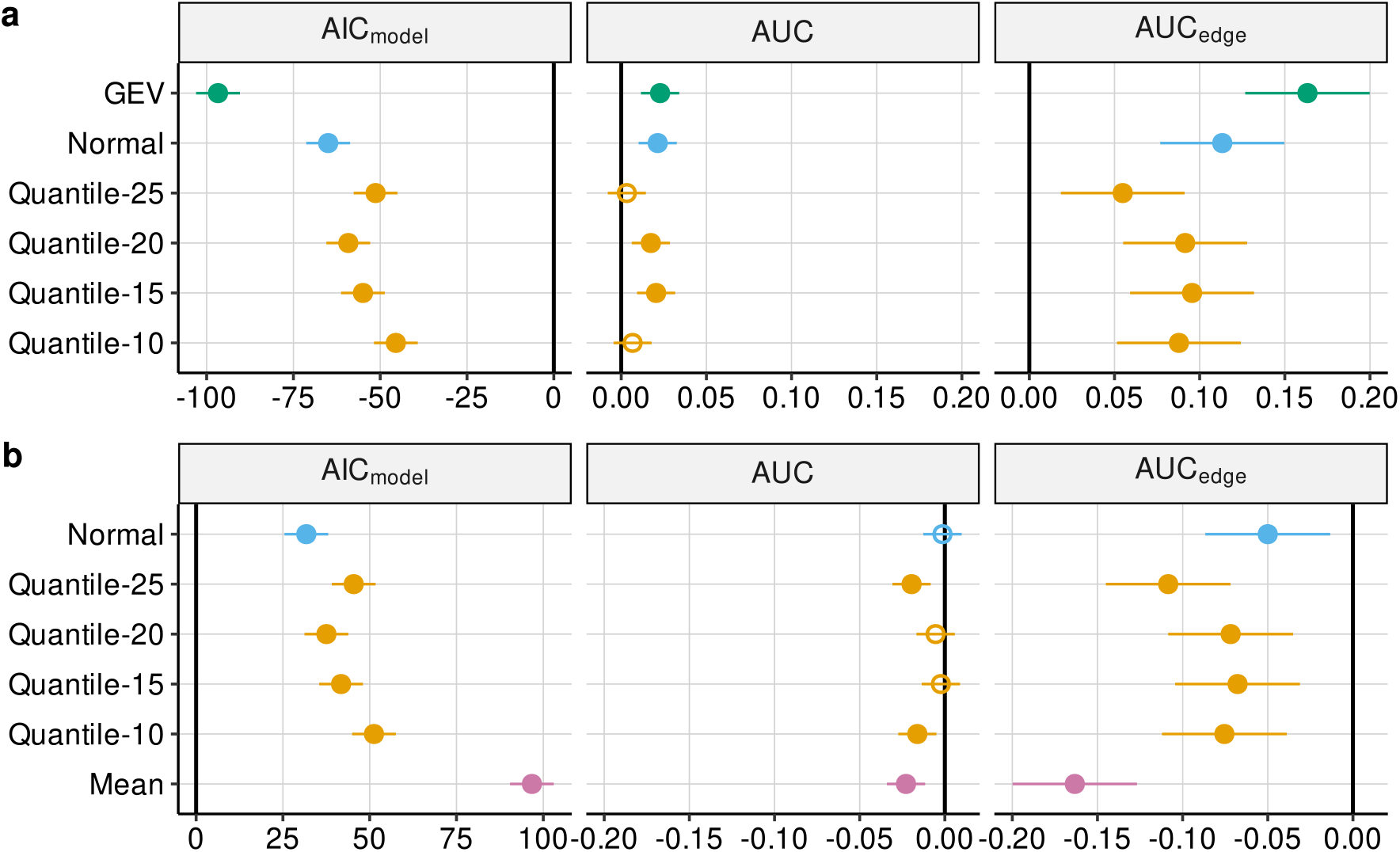
(a) Difference in AIC_model_, AUC and AUC_edge_ between the Mean model and the other modelling approaches and (b) difference in AIC_model_, AUC and AUC_edge_ between the GEV model and the other modelling approaches according to the mixed-effects models. The coloured lines represent the confidence interval with an α of 0.05. Significant differences compared to the reference model are shown using a filled point, while a hollow point represents the opposite.

Although the mixed-effects models allow us to derive general conclusions across all 28 modelled species, the difference in model fit and performance for each individual species varies (Appendix C).

### Spatial predictions

The spatial predictions by the Mean and GEV approaches describe a similar overall distribution for each species (Fig. 6 and Appendix I). Yet, considerable absolute differences arise in locations where strong influence by intense extremes can be expected (Fig. 6a). These are regions with considerable influence from the Atlantic Ocean, regions characterised by a strong continental climate and mountainous regions. The locations of these large absolute differences and their direction are highly species-dependent however. For example, the Pindus mountain range in Greece is considered less suitable by the GEV approach compared to the Mean approach for *Picea abies*, while it is considered more suitable for *Abies alba*. On the other hand, the relative differences show uniformly across species that the GEV approach decreases the predicted values at the distribution edges (Fig. 6b). For some species, such as *Picea abies*, this is also accompanied by a larger relative increase at the core of the species range.

**Figure 6.**
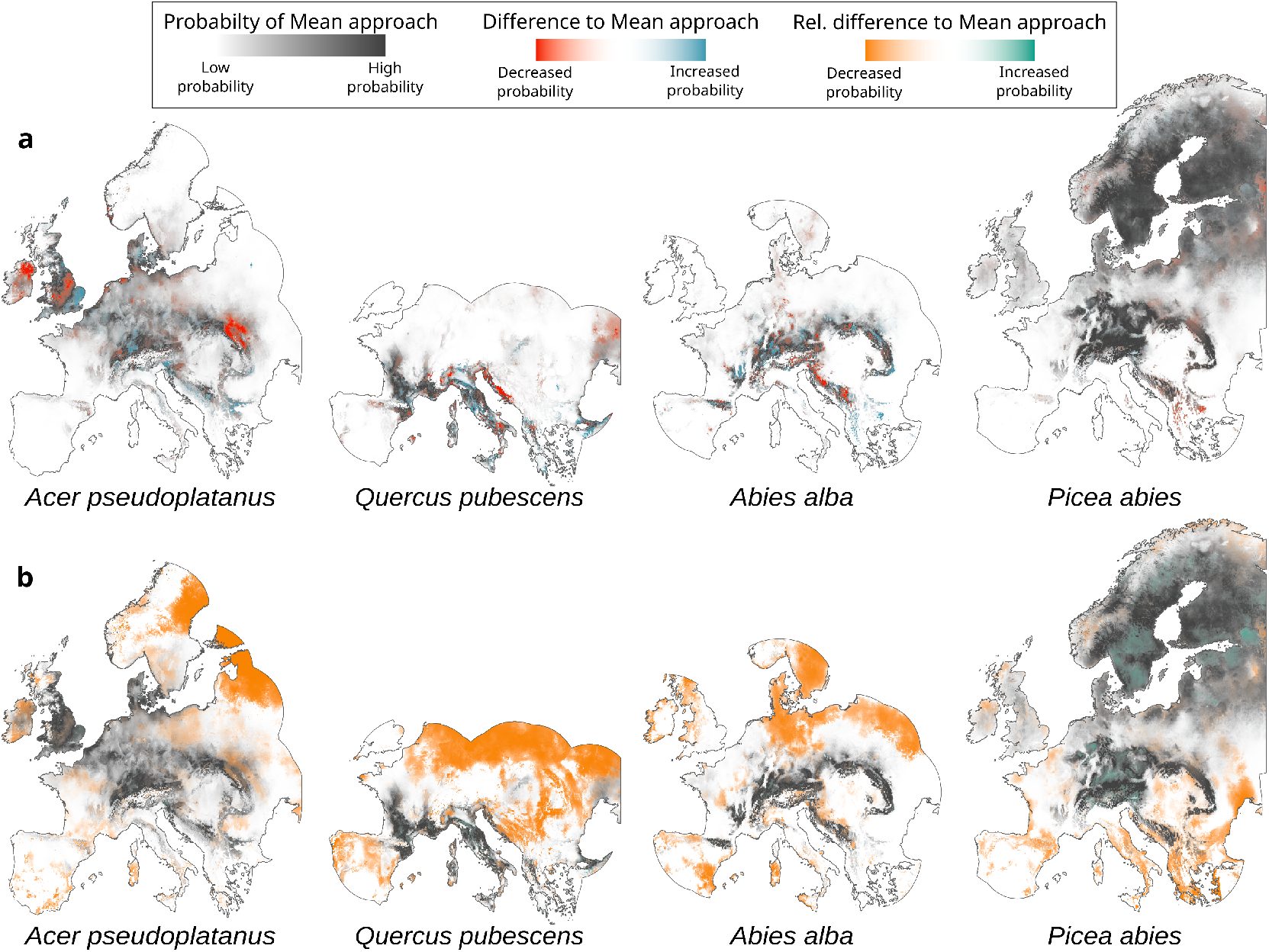
Spatial predictions based on the GEV approach are noticeably altered compared to those based on the Mean approach. A comparison between the spatial predictions derived from the GEV approach and the Mean approach are given for four tree species. Both the absolute difference as the relative difference are given. The probability for the species as predicted by the Mean approach is given in a gradient from white to dark grey (a and b), while the absolute difference in predicted probability between the GEV approach and the Mean approach is shown as an overlaid gradient from red to blue (a) and the relative difference in predicted probability between the GEV approach and the Mean approach is shown as an overlaid gradient from orange to green (b). More opaque red or orange indicates a greater decrease in probability while more opaque blue or green indicates a greater increase. Only the 500-km buffer around presences is shown. Predictions were performed on 1/24-degree spatial resolution.

## Discussion

Capturing more than solely the mean intensity of climatic extremes can improve the modelling of species distributions (Fig. 5), confirming the findings of Zimmermann et al. (2009) and Stewart et al. (2021). While these previous studies considered only models with a small set of predictor variables (a maximum of 6), we show this to be the case even when a large set of bioclimatic variables is considered in addition to extreme variables. The GEV approach outperforms all other approaches according to the AIC_model_, yet the cross-validated AUC shows a less cut-and-dry difference between the GEV, Normal and Quantile approaches. Given that the Mean models for the vast majority of species already have a high AUC, it is, however, no surprise that with such well-performing baseline, the observed differences in model AUC remain relatively small.

A major benefit of the GEV approach is that it complies to the theoretical expectations for the distribution of extreme variables. From the extreme value theory, we expect the extreme variables which were the focus of this analysis to follow the GEV distribution. The improved fit the GEV distribution provides over the normal distribution in almost all of Europe supports this theoretical underpinning (Fig. 4). In other words, capturing all combinations of frequency and intensity of these extremes while accounting for their skewed nature is relevant in the context of this analysis. Although empirical approaches, i.e. using mean, standard deviation, and empirical quantiles, cannot accurately capture all characteristics of extremes (Coles, 2001), they may still approximate these characteristics sufficiently, resulting in the similar overall model performance seen in our analysis. Similar-performing models do not necessarily predict similar distributions, however, especially when the models are projected to future climates which are far removed from the historical environment (Morán-Ordóñez et al., 2018). An area where the ability of the GEV approach to capture the skewed nature of extremes takes front stage is the distribution edges.

Species’ range margins, where conditions are overall less favourable to the species, are expected to be most influenced by extremes. Zimmermann et al. (2009) argued that asymmetric effects of severe extremes might create local sink populations which alter the margins otherwise predicted by climate means. The role that extremes play in these sink dynamics at the range edges of tree species is commonly identified (W. R. L. Anderegg et al., 2019; Sánchez-Salguero et al., 2017). The considerable improvement in AUC_edge_ indicates that the benefit of additional details of extreme variables incorporated in the SDMs manifests itself in distribution edges (Fig. 5). Moreover, the GEV approach outperforms all other models at the distribution edges, indicating that its superior ability to characterise the skewed distribution of extreme variables (Fig. 4) also allows it to better estimate the species edges. The spatial distribution of relative differences in predicted values between the GEV approach and the Mean approach shows a narrowing of the species range, with decreased estimates at the range edges (Fig. 6b).

The GEV approach, unlike any of the other approaches, provides explicit information on the behaviour of the intense tail-end of an extreme variable’s distribution in addition to its centre and spread characteristics. With climate change, this tail-end contains those extremes with most potential to alter species distribution early and abruptly (Zwiers et al., 2013; Turner et al., 2020). As demonstrated, a detailed characterisation of extreme variables is useful for the current conditions. This indicates that relevant differences in the distribution of extreme variables occur in centre, spread and skew across the study area. Not only differences in space exist, but changes over time in these three characteristics are also expected due to climate change (Coumou & Rahmstorf, 2012). In fact, changes in all three GEV parameters of the three extreme variables used in this analysis can already be observed under past climate change when comparing two historical 30-year periods, 1958-1987 and 1992-2021 (Appendix E). The trends in extremes do not necessarily follow changes in average climate conditions and therefore warrant separate in-depth consideration (Zwiers et al., 2013; Seneviratne et al., 2021). Consequently, the GEV approach can play a role in accurately projecting the effects of climate change. As Germain & Lutz (2020) point out, a more detailed characterisation of the environment also leads to more ways in which future environments can deviate from those in the training period. The choice of approach therefore determines what environments are considered novel in future scenarios. Novel environments, i.e. combinations of environmental conditions which are not present in the training data, pose a problem when projecting species distribution models. Projections to unsampled environments require extrapolation, which may be ecologically and statistically invalid (Fitzpatrick & Hargrove, 2009). The additional details on the distribution of extremes provided by the GEV distribution will allow differentiation between environments which previously were deemed alike. It is thus likely that incorporating extremes as proposed in this paper will reveal previously unaccounted-for extrapolation in model projections.

It has become abundantly clear that due to the changing climate, species are shifting their range (Lenoir et al., 2020). The ability of species to fully match the pace of climate change generates a need for effective conservation measures. Prioritising these efforts, especially when anticipating future climates, often (partly) relies on SDM outputs (Guisan et al., 2013). Assisted migration of species is one such application in conservation in which SDMs can play a crucial role to identify a need for such action and to identify potential recipient locations (Guisan et al., 2013). Prasad et al. (2020) identify a potential for near-range and extra-range assisted migration, which target areas just beyond or far beyond the current range limits respectively, to conserve forests under climate change. These types of assisted migration involve locations which currently are not or only marginally suitable for a species. Furthermore, the identification of climate refugia, areas with climatic conditions suitable for a species surrounded by a landscape which is not, is another critical application of SDMs in conservation (Baumgartner et al., 2018). For both applications, accurate SDM predictions are needed at the margins of a species’ niche, where the influence of extremes is most pronounced, and where the GEV approach improves upon the other approaches according to our results. The narrower niches predicted by the GEV approach could mean a more pessimistic outlook on which species are at risk and which locations are appropriate as recipients of assisted migration or as climate refugium.

For the projection of the GEV approach to future climate scenarios, monthly time series for those future scenarios are required. Although there are datasets that provide such data (e.g. Abatzoglou et al., 2018; Karger et al., 2020), the assumptions underlying these data are that the mean, or mean and standard deviation are of interest (e.g. Qin et al., 2020). As such, these datasets do not produce valid estimates for the GEV distribution parameters of extreme variables. In the case of the TerraClimate dataset (Abatzoglou et al., 2018), this translates to a shape parameter for temperature extremes which unrealistically shows no change between the historical and +4 °C future climate scenario in most of the study area (Appendix E). Other datasets widely used for species distribution modelling like WorldClim (Fick & Hijmans, 2017) and CHELSA (Karger et al., 2017), serve only the mean or the mean and the standard deviation for their future climatologies, even for extreme variables for which these metrics are not expected to fully describe the distribution. In addition, these sources do not provide access to the monthly data either, making it impossible to derive the GEV parameters from these data sources. For the European extent, an alternative is the output of regional climate models from the EURO-CORDEX initiative (Jacob et al., 2014). However, these may require bias correction to align model output with actual historical measurements, which also requires assumptions on distributions and moreover can affect extremes differently from means (Huang et al., 2014). The lack of data available for future climate scenarios currently hampers the implementation of this approach for climate change impact assessment. Since trends in the intensity and frequency of climatic extremes have been observed in many regions worldwide and are projected to continue in the future (Seneviratne et al., 2021), and changes in extremes are often the first to affect species (Zwiers et al., 2013; Turner et al., 2020), future climate data adequately capturing changes in all distributional parameters are urgently needed. We therefore encourage future climate scenario data providers to move beyond serving mean and standard deviation, and where feasible, provide time series on which further analysis can be conducted to realistically account for shifts in extreme climatic events.

Useful SDMs are first and foremost constructed from relevant environmental variables (Fourcade et al., 2018). Climatic extremes can pose a barrier to a species distribution and can therefore be ecologically relevant for modelling a species’ distribution. Whether climatic extremes are considered in detail should still depend on the ecological traits of the species of interest, and whether extremes are a focus of the analysis. If climatic extremes are considered in more detail in SDMs, the way these climatic extremes are characterised should also strive towards maximising relevance. As shown in this paper, this translates into using the location, scale, and shape parameters of the GEV distribution of climatic extreme variables that arise following the extreme value theory. The optimal characterisation of extreme variables through these parameters of the GEV distribution allows for a better characterisation of species distributions and especially their edges.

## Data accessibility statement

The code and data contained in this paper are available in a GitHub repository (link anonymised).

