## Appendix for "Incorporating climatic extremes using the GEV distribution improves SDM range edge performance"

### Appendix A

#### Bioclimatic variables

The description of the 19 bioclimatic variables used in this analysis is given here (Fick & Hijmans, 2017):

- Bio1 = Annual Mean Temperature
- Bio2 = Mean Diurnal Range (Mean of monthly ($T_{max}$ - $T_{min}$))
- Bio3 = Isothermality (Bio2/Bio7) ($\times$100)
- Bio4 = Temperature Seasonality (standard deviation $\times$100)
- Bio5 = Max Temperature of Warmest Month
- Bio6 = Min Temperature of Coldest Month
- Bio7 = Temperature Annual Range (Bio5-Bio6)
- Bio8 = Mean Temperature of Wettest Quarter
- Bio9 = Mean Temperature of Driest Quarter
- Bio10 = Mean Temperature of Warmest Quarter
- Bio11 = Mean Temperature of Coldest Quarter
- Bio12 = Annual Precipitation
- Bio13 = Precipitation of Wettest Month
- Bio14 = Precipitation of Driest Month
- Bio15 = Precipitation Seasonality (Coefficient of Variation)
- Bio16 = Precipitation of Wettest Quarter
- Bio17 = Precipitation of Driest Quarter
- Bio18 = Precipitation of Warmest Quarter
- Bio19 = Precipitation of Coldest Quarter

### Appendix B

#### Species records

##### Table B1

*The number of presence and absence records per species after filtering to the calibration area of 500 km around presences and aggregating to the 1/24-degree resolution of the climate data. Additionally, the range of cross-validation (CV) partitions and the range of presences and absences present in each partition is given for each species.*

| Species | Presences | Absences | Range of number of CV partitions | Range of CV partition presences | Range of CV partition absences |
|---|---|---|---|---|---|
| Abies alba | 6,709 | 80,540 | 5-6 | 89-192 | 1222-1778 |
| Acer campestre | 5,009 | 91,479 | 4-5 | 97-169 | 1552-3716 |
| Acer pseudoplatanus | 10,401 | 98,251 | 7-8 | 109-215 | 1100-1637 |
| Alnus glutinosa | 8,749 | 113,492 | 6-6 | 123-172 | 1540-2102 |
| Alnus incana | 5,971 | 101,218 | 4-5 | 108-201 | 1922-2845 |
| Betula pendula | 16,990 | 105,806 | 10-10 | 132-209 | 892-1278 |
| Betula pubescens | 19,952 | 103,069 | 10-10 | 126-295 | 866-1219 |
| Carpinus betulus | 10,677 | 93,291 | 7-9 | 104-241 | 731-1823 |
| Castanea sativa | 6,653 | 81,501 | 6-6 | 89-150 | 1183-1590 |
| Fagus sylvatica | 24,132 | 82,070 | 10-10 | 109-446 | 413-1261 |
| Fraxinus excelsior | 14,608 | 108,794 | 9-10 | 125-189 | 939-1306 |
| Larix decidua | 6,871 | 107,561 | 4-5 | 115-256 | 1902-3399 |
| Picea abies | 46,607 | 73,250 | 10-10 | 154-928 | 232-1344 |
| Picea sitchensis | 4,868 | 76,852 | 4-5 | 83-166 | 991-2436 |
| Pinus halepensis | 4,736 | 50,683 | 5-5 | 67-126 | 816-1124 |
| Pinus nigra | 5,091 | 91,004 | 4-4 | 98-167 | 2020-2581 |
| Pinus pinaster | 7,389 | 65,982 | 7-7 | 76-150 | 745-1162 |
| Pinus sylvestris | 46,358 | 77,102 | 10-10 | 162-909 | 331-1338 |
| Populus tremula | 9,486 | 113,956 | 6-7 | 124-215 | 1452-2028 |
| Prunus avium | 5,305 | 102,994 | 4-4 | 109-167 | 1494-3215 |
| Pseudotsuga menziesii | 4,332 | 91,985 | 4-4 | 97-135 | 1392-2954 |
| Quercus ilex | 10,468 | 58,550 | 10-10 | 72-182 | 444-758 |
| Quercus petraea | 12,931 | 84,381 | 9-10 | 98-174 | 732-1230 |
| Quercus pubescens | 5,895 | 66,934 | 5-6 | 73-187 | 887-1715 |
| Quercus robur | 22,975 | 94,695 | 10-10 | 127-427 | 708-1293 |
| Robinia pseudoacacia | 4,160 | 89,619 | 3-4 | 94-183 | 1434-4048 |
| Salix caprea | 6,379 | 117,059 | 4-4 | 124-212 | 2233-3776 |
| Sorbus aucuparia | 9,961 | 113,481 | 6-7 | 124-215 | 1429-2024 |

### Appendix C

#### Model fit and performance: tables

##### Table C1

*Average $AIC_{model}$ values for the models of the Mean, GEV, Normal and Quantile approaches*

| Species | GEV model | Normal model | Quantile-10 model | Quantile-15 model | Quantile-20 model | Quantile-25 model | Mean model |
|---|---|---|---|---|---|---|---|
| Abies alba | 2705 | 2722 | 2788 | 2792 | 2790 | 2797 | 2815 |
| Acer campestre | 3012 | 3040 | 3074 | 3054 | 3046 | 3059 | 3112 |
| Acer pseudoplatanus | 5022 | 5049 | 5078 | 5089 | 5091 | 5094 | 5132 |
| Alnus glutinosa | 5140 | 5155 | 5180 | 5185 | 5177 | 5176 | 5206 |
| Alnus incana | 3351 | 3409 | 3407 | 3391 | 3390 | 3401 | 3425 |
| Betula pendula | 7259 | 7301 | 7343 | 7305 | 7320 | 7305 | 7376 |
| Betula pubescens | 4439 | 4509 | 4526 | 4513 | 4511 | 4524 | 4593 |
| Carpinus betulus | 4215 | 4252 | 4288 | 4233 | 4243 | 4262 | 4343 |
| Castanea sativa | 2972 | 3019 | 3035 | 3053 | 3046 | 3053 | 3085 |
| Fagus sylvatica | 7072 | 7132 | 7178 | 7156 | 7137 | 7160 | 7275 |
| Fraxinus excelsior | 6619 | 6663 | 6704 | 6672 | 6672 | 6679 | 6744 |
| Larix decidua | 3202 | 3243 | 3218 | 3232 | 3231 | 3231 | 3270 |
| Picea abies | 8208 | 8292 | 8291 | 8264 | 8249 | 8265 | 8436 |
| Picea sitchensis | 1348 | 1367 | 1359 | 1353 | 1352 | 1353 | 1375 |
| Pinus halepensis | 1102 | 1121 | 1119 | 1119 | 1133 | 1131 | 1134 |
| Pinus nigra | 2626 | 2654 | 2666 | 2666 | 2666 | 2669 | 2686 |
| Pinus pinaster | 2379 | 2412 | 2419 | 2418 | 2404 | 2411 | 2450 |
| Pinus sylvestris | 10702 | 10664 | 10759 | 10684 | 10709 | 10756 | 10905 |
| Populus tremula | 5617 | 5663 | 5652 | 5645 | 5653 | 5657 | 5688 |
| Prunus avium | 3478 | 3481 | 3512 | 3490 | 3485 | 3494 | 3532 |
| Pseudotsuga menziesii | 2740 | 2739 | 2756 | 2761 | 2730 | 2742 | 2785 |
| Quercus ilex | 2650 | 2669 | 2682 | 2676 | 2650 | 2661 | 2700 |
| Quercus petraea | 4961 | 5017 | 5018 | 5011 | 5004 | 5012 | 5059 |
| Quercus pubescens | 2001 | 2029 | 2023 | 2043 | 2045 | 2059 | 2104 |
| Quercus robur | 7565 | 7618 | 7634 | 7676 | 7645 | 7648 | 7714 |
| Robinia pseudoacacia | 2104 | 2103 | 2112 | 2101 | 2101 | 2097 | 2115 |
| Salix caprea | 4476 | 4504 | 4511 | 4490 | 4502 | 4512 | 4538 |
| Sorbus aucuparia | 5574 | 5599 | 5641 | 5638 | 5608 | 5601 | 5649 |

**Table C2**

*Average AUC values for the models of the Mean, GEV, Normal and Quantile approaches*

| Species | Mean model | GEV model | Normal model | Quantile-10 model | Quantile-15 model | Quantile-20 model | Quantile-25 model |
|---|---|---|---|---|---|---|---|
| Abies alba | 0.909 | 0.915 | 0.916 | 0.911 | 0.910 | 0.909 | 0.908 |
| Acer campestre | 0.744 | 0.754 | 0.753 | 0.745 | 0.752 | 0.752 | 0.748 |
| Acer pseudoplatanus | 0.829 | 0.838 | 0.836 | 0.834 | 0.832 | 0.829 | 0.830 |
| Alnus glutinosa | 0.776 | 0.778 | 0.780 | 0.775 | 0.776 | 0.775 | 0.774 |
| Alnus incana | 0.828 | 0.824 | 0.820 | 0.828 | 0.824 | 0.825 | 0.822 |
| Betula pendula | 0.822 | 0.826 | 0.824 | 0.821 | 0.827 | 0.824 | 0.825 |
| Betula pubescens | 0.950 | 0.953 | 0.952 | 0.951 | 0.952 | 0.952 | 0.951 |
| Carpinus betulus | 0.888 | 0.894 | 0.892 | 0.890 | 0.895 | 0.892 | 0.891 |
| Castanea sativa | 0.874 | 0.877 | 0.879 | 0.878 | 0.876 | 0.875 | 0.873 |
| Fagus sylvatica | 0.836 | 0.839 | 0.839 | 0.833 | 0.841 | 0.840 | 0.838 |
| Fraxinus excelsior | 0.825 | 0.832 | 0.831 | 0.826 | 0.830 | 0.829 | 0.829 |
| Larix decidua | 0.900 | 0.899 | 0.898 | 0.905 | 0.902 | 0.900 | 0.901 |
| Picea abies | 0.872 | 0.877 | 0.873 | 0.877 | 0.878 | 0.878 | 0.877 |
| Picea sitchensis | 0.968 | 0.968 | 0.965 | 0.968 | 0.968 | 0.967 | 0.968 |
| Pinus halepensis | 0.958 | 0.951 | 0.952 | 0.956 | 0.953 | 0.953 | 0.952 |
| Pinus nigra | 0.816 | 0.813 | 0.820 | 0.813 | 0.819 | 0.822 | 0.818 |
| Pinus pinaster | 0.895 | 0.886 | 0.890 | 0.891 | 0.895 | 0.897 | 0.896 |
| Pinus sylvestris | 0.771 | 0.771 | 0.781 | 0.773 | 0.783 | 0.778 | 0.773 |
| Populus tremula | 0.765 | 0.770 | 0.765 | 0.768 | 0.769 | 0.766 | 0.767 |
| Prunus avium | 0.754 | 0.758 | 0.769 | 0.751 | 0.767 | 0.763 | 0.754 |
| Pseudotsuga menziesii | 0.786 | 0.781 | 0.787 | 0.783 | 0.784 | 0.794 | 0.790 |
| Quercus ilex | 0.932 | 0.932 | 0.933 | 0.933 | 0.933 | 0.933 | 0.933 |
| Quercus petraea | 0.870 | 0.870 | 0.869 | 0.870 | 0.871 | 0.871 | 0.870 |
| Quercus pubescens | 0.912 | 0.921 | 0.916 | 0.921 | 0.918 | 0.916 | 0.912 |
| Quercus robur | 0.829 | 0.834 | 0.832 | 0.830 | 0.825 | 0.829 | 0.828 |
| Robinia pseudoacacia | 0.859 | 0.856 | 0.858 | 0.858 | 0.857 | 0.854 | 0.851 |
| Salix caprea | 0.632 | 0.659 | 0.645 | 0.638 | 0.638 | 0.637 | 0.633 |
| Sorbus aucuparia | 0.801 | 0.807 | 0.805 | 0.800 | 0.801 | 0.805 | 0.806 |

**Table C3**

*Average $AUC_{edge}$ values for the models of the Mean, GEV, Normal and Quantile approaches*

| Species | Mean model | GEV model | Normal model | Quantile-10 model | Quantile-15 model | Quantile-20 model | Quantile-25 model |
|---|---|---|---|---|---|---|---|
| Abies alba | 0.590 | 0.687 | 0.701 | 0.645 | 0.611 | 0.605 | 0.592 |
| Acer campestre | 0.531 | 0.608 | 0.594 | 0.576 | 0.600 | 0.577 | 0.553 |
| Acer pseudoplatanus | 0.617 | 0.671 | 0.659 | 0.661 | 0.650 | 0.626 | 0.634 |
| Alnus glutinosa | 0.568 | 0.586 | 0.591 | 0.571 | 0.576 | 0.567 | 0.567 |
| Alnus incana | 0.536 | 0.603 | 0.536 | 0.560 | 0.541 | 0.575 | 0.572 |
| Betula pendula | 0.608 | 0.638 | 0.612 | 0.613 | 0.633 | 0.620 | 0.633 |
| Betula pubescens | 0.727 | 0.820 | 0.789 | 0.769 | 0.783 | 0.764 | 0.765 |
| Carpinus betulus | 0.593 | 0.654 | 0.626 | 0.605 | 0.632 | 0.608 | 0.597 |
| Castanea sativa | 0.588 | 0.670 | 0.659 | 0.649 | 0.626 | 0.616 | 0.600 |
| Fagus sylvatica | 0.667 | 0.690 | 0.703 | 0.690 | 0.694 | 0.694 | 0.681 |
| Fraxinus excelsior | 0.606 | 0.627 | 0.624 | 0.595 | 0.614 | 0.609 | 0.606 |
| Larix decidua | 0.553 | 0.600 | 0.563 | 0.623 | 0.596 | 0.592 | 0.584 |
| Picea abies | 0.659 | 0.689 | 0.668 | 0.689 | 0.700 | 0.693 | 0.688 |
| Picea sitchensis | 0.583 | 0.645 | 0.631 | 0.585 | 0.640 | 0.632 | 0.627 |
| Pinus halepensis | 0.586 | 0.549 | 0.548 | 0.537 | 0.581 | 0.572 | 0.535 |
| Pinus nigra | 0.534 | 0.551 | 0.557 | 0.555 | 0.557 | 0.570 | 0.546 |
| Pinus pinaster | 0.585 | 0.561 | 0.544 | 0.566 | 0.593 | 0.598 | 0.595 |
| Pinus sylvestris | 0.629 | 0.618 | 0.656 | 0.636 | 0.657 | 0.641 | 0.627 |
| Populus tremula | 0.562 | 0.574 | 0.568 | 0.568 | 0.568 | 0.565 | 0.566 |
| Prunus avium | 0.560 | 0.595 | 0.625 | 0.561 | 0.607 | 0.604 | 0.588 |
| Pseudotsuga menziesii | 0.534 | 0.590 | 0.571 | 0.567 | 0.538 | 0.589 | 0.578 |
| Quercus ilex | 0.646 | 0.677 | 0.673 | 0.676 | 0.656 | 0.647 | 0.655 |
| Quercus petraea | 0.622 | 0.612 | 0.591 | 0.599 | 0.610 | 0.607 | 0.604 |
| Quercus pubescens | 0.609 | 0.696 | 0.663 | 0.691 | 0.657 | 0.656 | 0.630 |
| Quercus robur | 0.654 | 0.698 | 0.689 | 0.672 | 0.637 | 0.659 | 0.641 |
| Robinia pseudoacacia | 0.550 | 0.584 | 0.549 | 0.599 | 0.562 | 0.580 | 0.557 |
| Salix caprea | 0.523 | 0.549 | 0.559 | 0.550 | 0.537 | 0.538 | 0.532 |
| Sorbus aucuparia | 0.604 | 0.649 | 0.620 | 0.597 | 0.600 | 0.627 | 0.634 |

### Appendix D

#### Model fit and performance: boxplot graphs

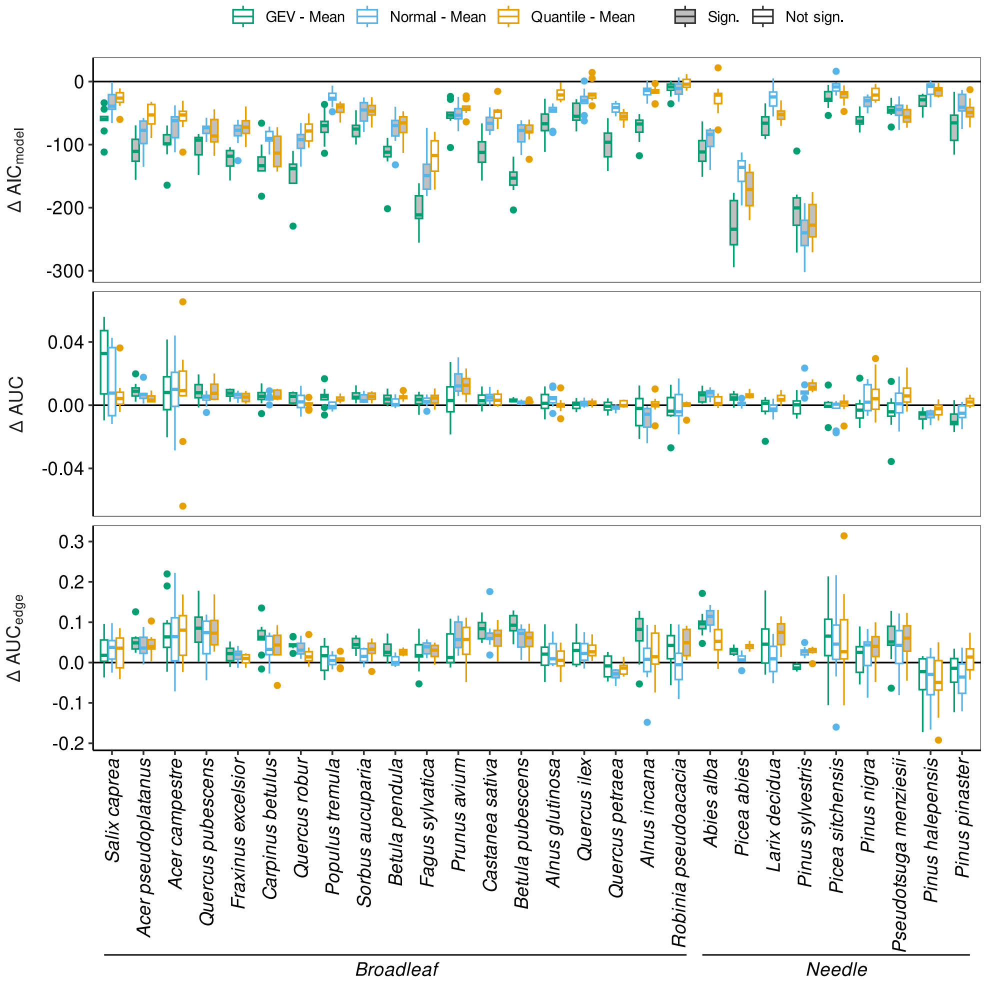


**Figure D1**

*The difference in $AIC_{model}$, $AUC$ and $AUC_{edge}$ for the GEV, Normal, and Quantile models compared to the Mean model is given. Per species, only one Quantile model which yields the best AUC across the tend random folds is given. Boxplots represent values from the 10 random folds. Species are grouped into leaf type and ordered within groups by decreasing $\Delta AUC$ between the GEV and Mean model. Filled boxplots indicate a significant difference compared to the model approach according to the Friedman test and Nemenyi's many-to-one post-hoc test.*

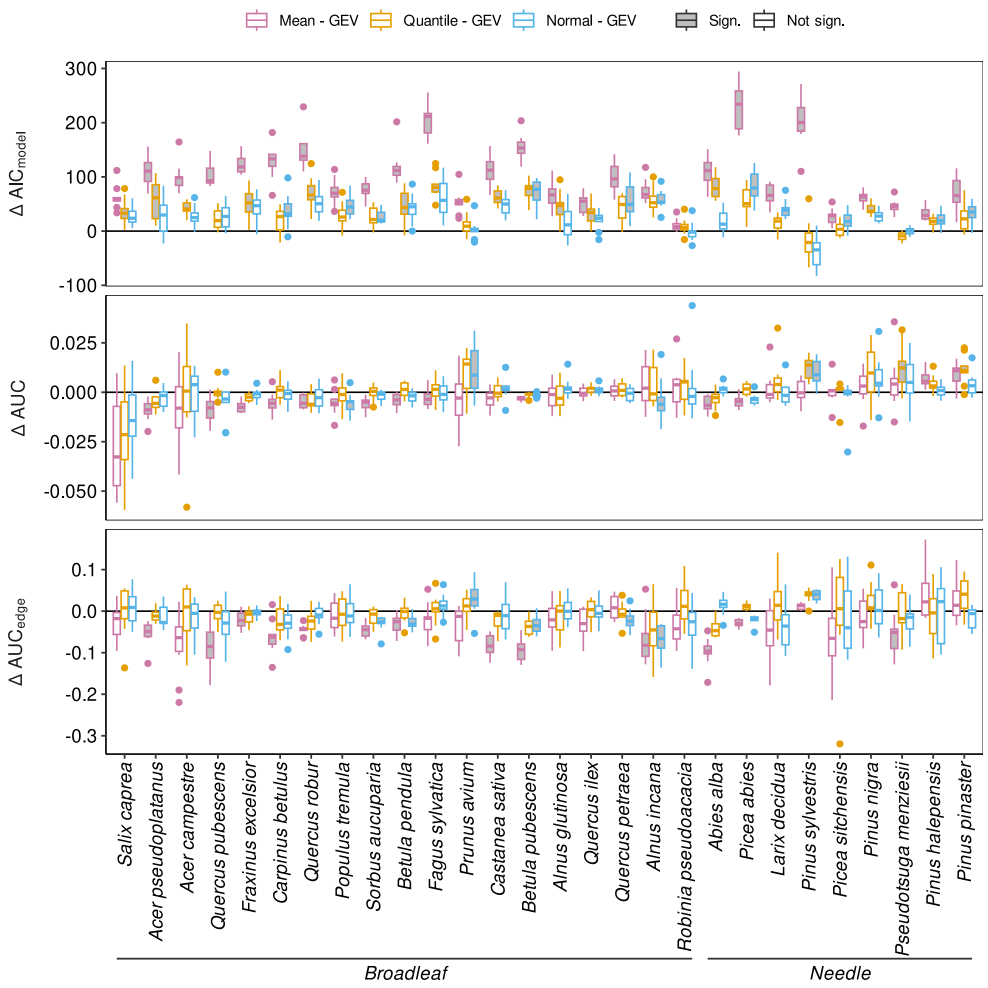


731 **Figure D2**

*The difference in $AIC_{model}$, $AUC$ and $AUC_{edge}$ for the Mean, Normal and Quantile models compared to the GEV model is given. Per species, only one Quantile model which yields the best AUC across the tend random folds is given. Boxplots represent values from each of the 10 random folds. Species are grouped into leaf type and ordered within groups by decreasing $\Delta AUC$ between the GEV and Mean model. Filled boxplots indicate a significant difference compared to the reference model according to the*

732 *Friedman test and Nemenyi's many-to-one post-hoc test*

### Appendix E

#### Changes in GEV parameter estimates

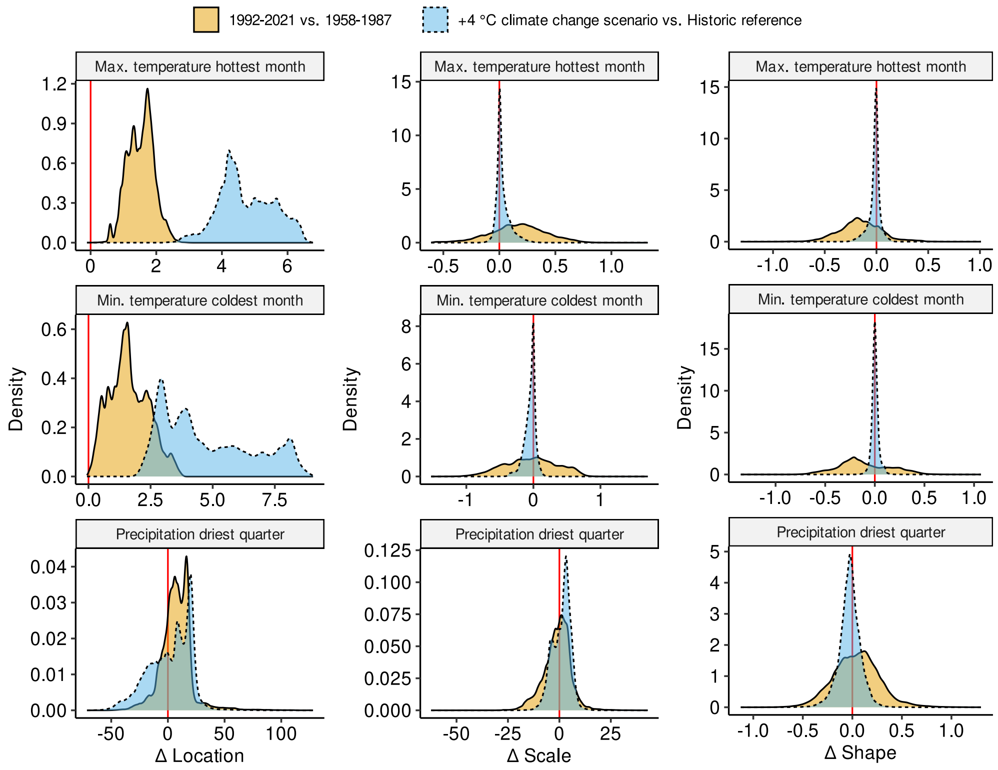


733 **Figure E1**

*Density distribution of the difference in GEV distribution parameters for the maximum temperature of the hottest month (TxHm), the minimum temperature of the coldest month (TnCm) and the rolling quarterly precipitation of the driest quarter (PqDq) for two comparisons: 1992-2021 compared to 1958-1987, and the +4 °C TerraClimate climate change scenario compared to the historic references period (1986-2015) for all*
734 *pixels in the study area.*

### Appendix F

#### Correlation matrix of climatic extreme variables

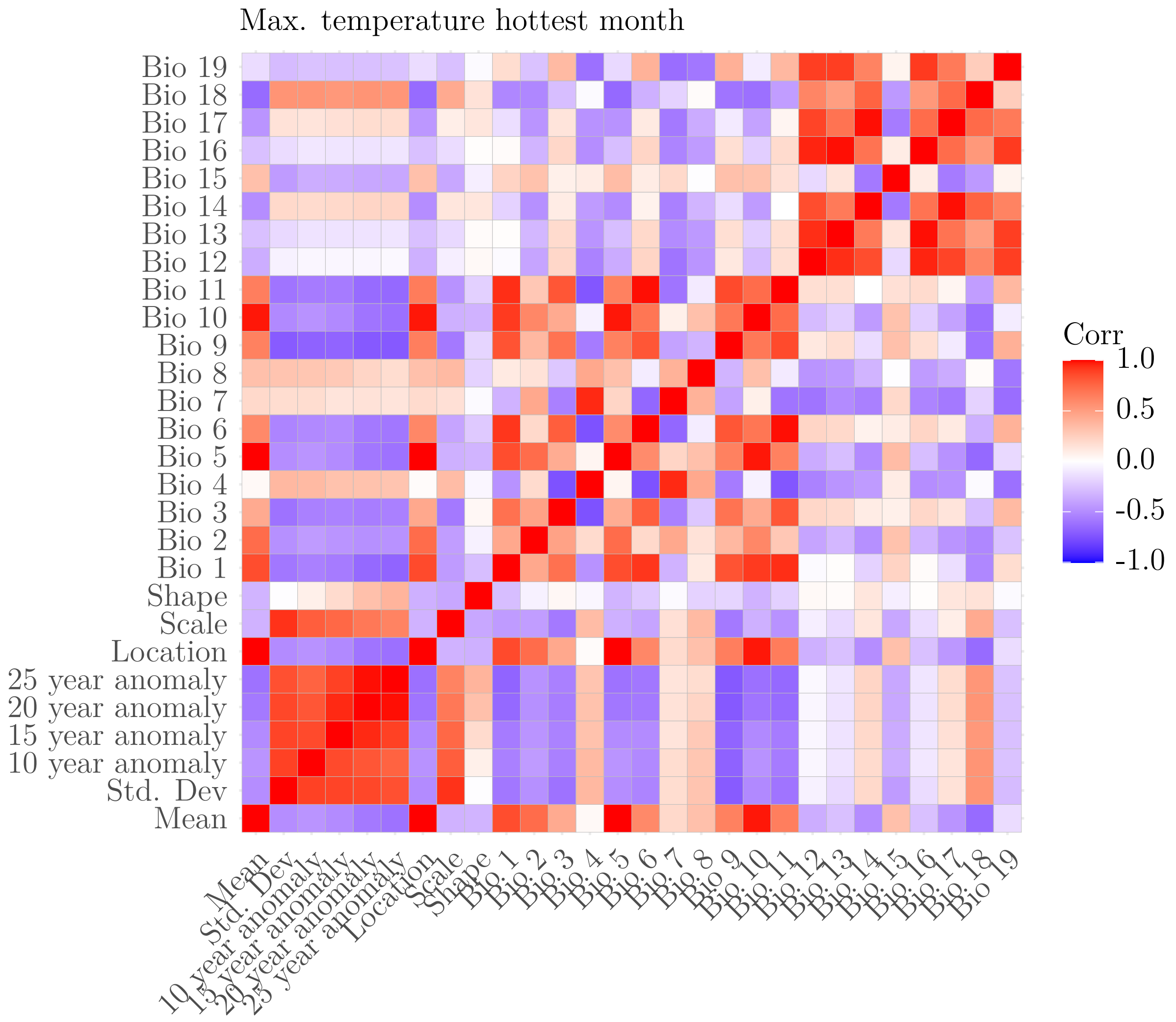


**Figure F1**

*Visualisation of the correlation matrix of the different parameters derived from the maximum temperature of the hottest month and the 19 bioclimatic variables. The 10-, 15-, 20- and 25-year anomalies are the difference of the extreme intensities for the corresponding return period and the mean.*

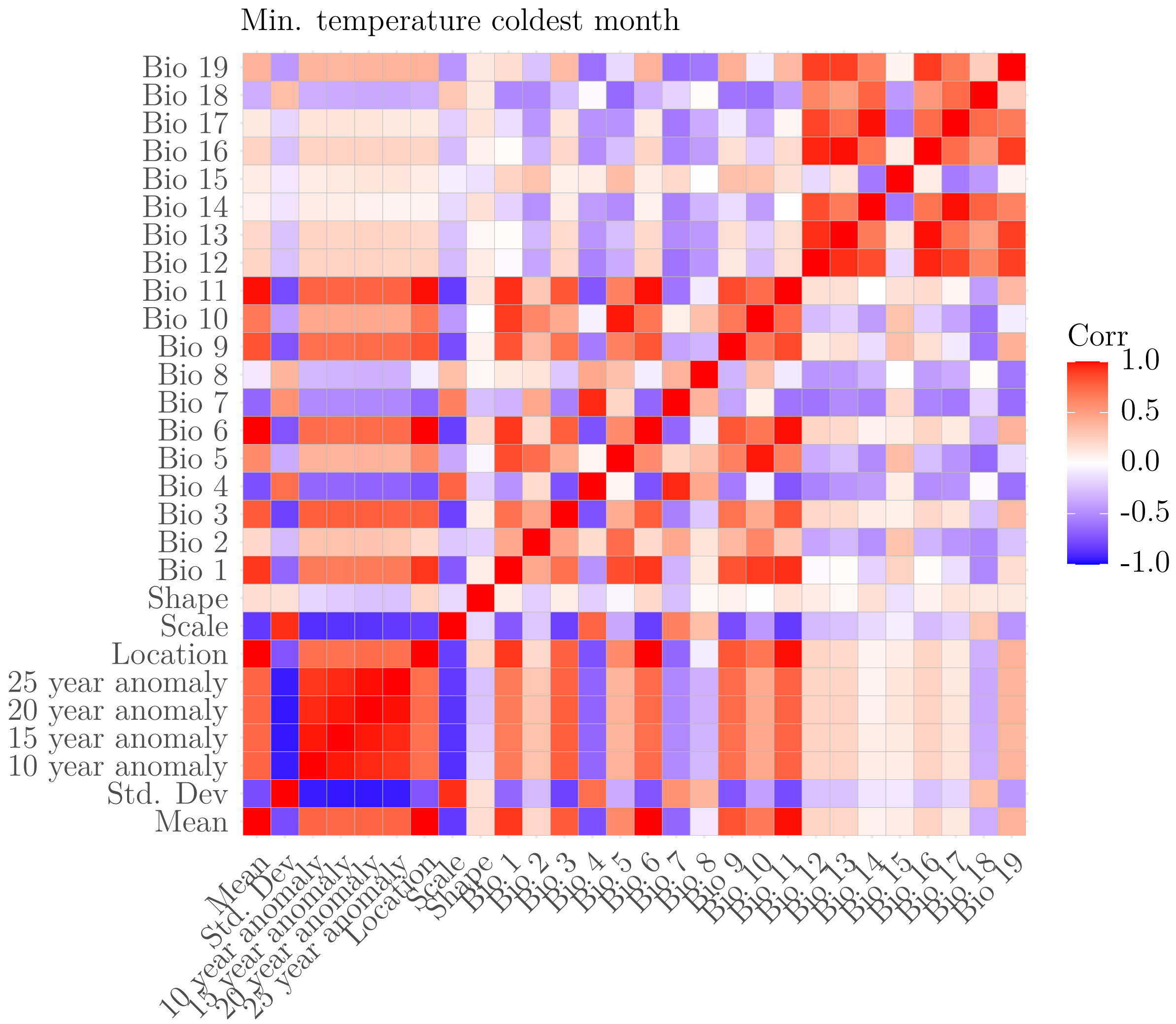


737 **Figure F2**

*Visualisation of the correlation matrix of the different parameters derived from the minimum temperature of the coldest month and the 19 bioclimatic variables. The 10-, 15-, 20- and 25-year anomalies are the difference of the extreme intensities for the*

738 *corresponding return period and the mean.*

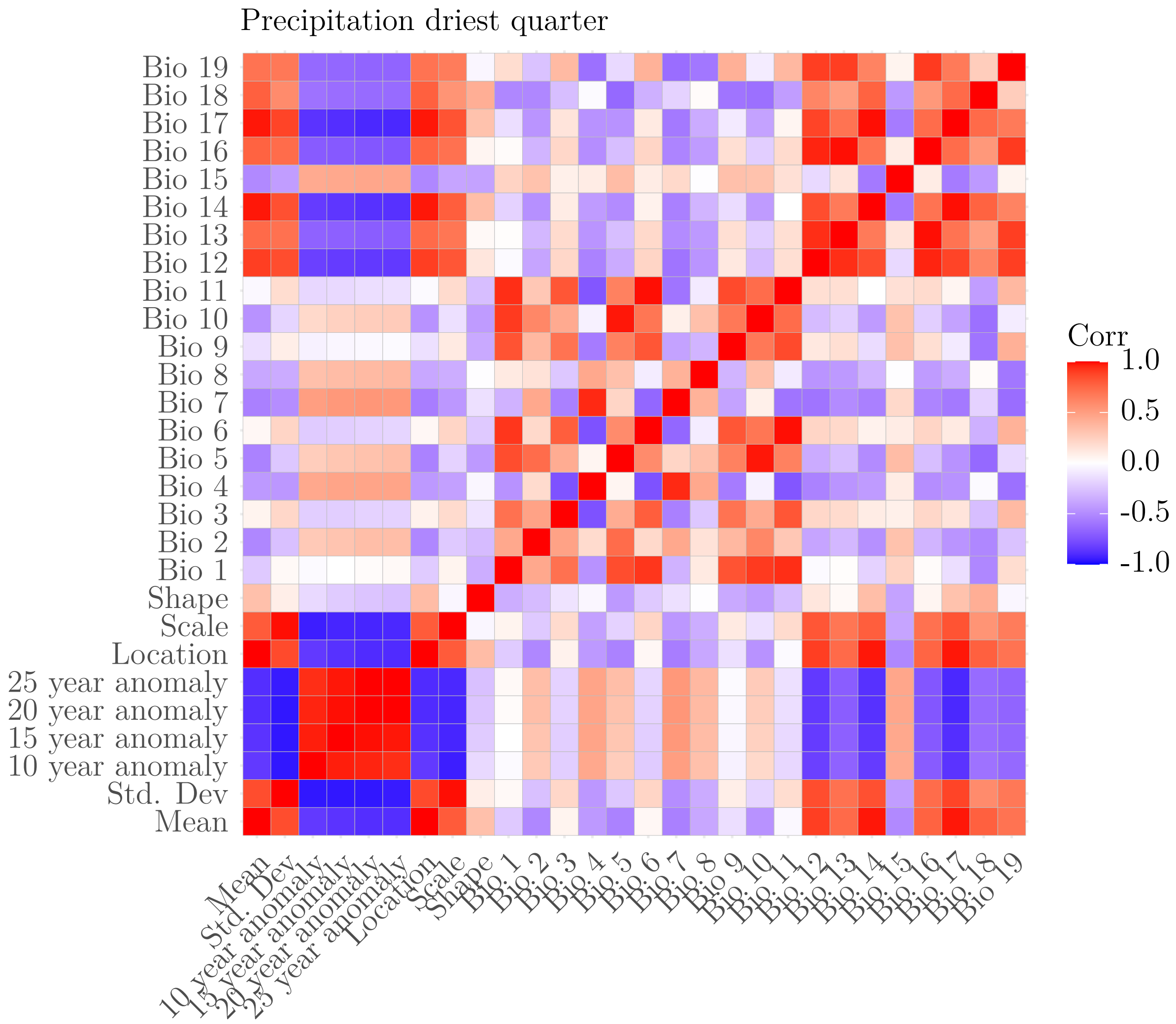


739 **Figure F3**

*Visualisation of the correlation matrix of the different parameters derived from the precipitation of the driest quarter and the 19 bioclimatic variables. The 10-, 15-, 20- and 25-year anomalies are the difference of the extreme intensities for the*

740 *corresponding return period and the mean.*

### Appendix G

#### ODMAP

### ODMAP

| | | |
|---|---|---|
| 741 | Authorship | · Authors: Anonymised |
|  |  | · Contact email: Anonymised |
|  |  | · Title: Incorporating climatic extremes using the GEV distribution improves SDM range edge performance |
| | Model objective | · SDM objective/purpose: explanation, comparison of the effect of modelling approach on model performance |
| | | · Main target output: continuous suitability index for each species. |
| | Taxon | · 28 European tree species with differing ecological traits |
| | Location | Europe up to a longitude of 34° East, the British Isles, and Mediterranean islands excluding Malta and Cyprus. |
| | Scale of analysis | · Spatial Extent (Lon / Lat) 12.3° W to 34° E, 34.5° N to 74.4° N |
| | | · Spatial resolution: 1/24-degree |
| | | · Temporal extent/time period: 1986 until 2015 |
| | | · Temporal resolution, if applicable: One period |
| | | · Type of extent boundary: natural |
| | Biodiversity data overview | · Observation type: Standardised monitoring data and field survey |
| | | · Response/data type: presence-absence |
| **O**verview | Type of predictors | Climatic |
| | Conceptual model | · We tested whether modelling species distrubution using the generalised extreme value distribution parameters derived from extreme variables yielded better model fit, performance and spatial predictions compared to models using mean, mean and standard deviation, and mean and return level for a set of return periods. Apart from the predictor variables derived from the extreme variables, an additional 16 bioclimatic variabels are included in the models. |
| | Assumptions | · We assumed that: there is a species-environment equilibrium, all plots where a species was not recorded to be an absence, no spatial bias remained in the processed occurrence dataset, the extreme variables included in the model can pose a barrier to the species' distribution through acute stress. |
| | SDM algorithms | · Model algorithms: Generalised Addititve Models (GAMs) |
|  |  | · Justification of model complexity: To aid comparability across models, a linear combination of smooth terms was used. We fixed the paramter setting for the smooth terms of k = 10 following Valavi et al. (2021). |
| | | · Ensembles: an unweighted mean ensemble was made for combination of species and approach to make spatial predictions |
|  | Model workflow | · Conceptual description of modelling steps including model fitting, assessment and prediction |
| | Software, codes and data | · Software: All analysis was conducted usig R version 4.2.2 (R Core Team, 2022) with the *flexsdm* (Velazco et al., 2022) , *mgcv* (Wood, 2011) and *dismo* (Hijmans, Phillips, & Elith, 2022) packages being instrumental to modelling. |
| | | · Code and Data availability: Github repositroy. See data archiving statement on publication |
|  | Biodiversity data | · Taxon names: *Abies alba*, *Acer campestre*, *Acer pseudoplatanus*, *Alnus glutinosa*, *Alnus incana*, *Betula pendula*, *Betula pubescens*, *Carpinus betulus*, *Castanea sativa*, *Fagus sylvatica*, *Fraxinus excelsior*, *Larix decidua*, *Picea abies*, *Picea sitchensis*, *Pinus halepensis*, *Pinus nigra*, *Pinus pinaster*, *Pinus sylvestris*, *Populus tremula*, *Prunus avium*, *Pseudotsuga menziesii*, *Quercus ilex*, *Quercus petraea*, *Quercus pubescens*, *Quercus robur*, *Robinia pseudoacacia*, *Salix caprea and Sorbus aucuparia* |
| | | · Details on taxonomic reference system: See Mauri *et al.*, 2017 |
| | | · Ecological level: Species |
| | | · Biodiversity data source: EU-Forest aggregated forest inventory and research network data |
| | | · Sampling design: spatial design, separate per source in the EU-forest dataset |
|  |  | · Sample size per taxon: presences ranging from 4,160 to 46,607 and absences from 50,683 to 117,059. |
| | | · Details on scaling: The occurrence records were spatially thinned to the 1/24-degree resolution |
| **D**ata | | · Details on absence data collection: As only widely recognised species were used, any plot where a species was not noted as present was taken as an absence. Absences for a species were only kept if they were within 500 km from a presence of that species. |
| | | · Details on background data derivation, if applicable: / |
| | Data partitioning | 10 random folds, within each of which a spatially blocked cross validation was performed. |
| | Predictor variables | · State predictor variables used: Various parameters derived from maximum temperature of the hottest month, minimum temperature of the coldest month and precipitation of the driest quarter. Additionally 16 bioclimatic variables were included (all 19 biocliamtic variables excluding bio5, bio6 and bio17). |
| | | · Details on data sources: Monthly timeseries of maximum temperature, minimum temperature and precipitation from the TerraClimate dataset (Abatzoglou *et al.*, 2018). |
| | | · Spatial resolution and spatial extent of raw data: identical spatial extent, 1/24 degree resolution |
| | | · Map projection: WGS 84 |
| | | · Temporal resolution and temporal extent of raw data: monthly timeseries from 1986 to 2015 |
| | Variable pre-selection | · Details on pre-selection of variables: variables representing climatic extremes and 16 bioclimatic variables |
| | Multicollinearity | · Multicollinearity: to ensure the only differences between approaches were which predictor variables derived from extreme variables were included in the models, no action was taken regarding multicollinearity. This maximises the comparability between approaches. Additionally, as the analysis only concerns prediction rather than model parameter interpretation, multicollinearity does not pose a problem. |
| | Model settings | · Models settings for all selected algorithms: smooth terms as penalised regression splines with 10 basis functions and smoothing parameter estimation using the generalised cross validation criterion. Model terms were not penalised. |
| | | · Extrapolation beyond sample range: No |
| **M**odel | Model estimates | Coefficients on their own were not analysed in this analysis |

| | | |
|---|---|---|
| Model | Model selection / Model averaging / Ensembles | · Ensemble method: an unweighted mean ensemble was made for combination of species and approach to make spatial predictions |
| | Non-independence correction/analyses | None |
| | Threshold selection | True skill statistic (TSS) maximising threshold |
| **A**ssessment | Performance statistics | Area under the operating characteristic curve were calculated on validation data following a spatially blocked cross-validation. |
| | Plausibility check | Map display |
| **P**rediction | Prediction output | · Prediction unit: continuous suitability index for each species<br>· Visualisation/treatment of novel environments: masking |

### Appendix H

#### Feng *et al.* 2019 checklist

| Workflow | Category | What to report | Method and rationale |
|---|---|---|---|
| (A) Obtaining and processing occurrence data | metadata | (A1) source of occurrence data | EU-Forest (Mauri *et al., 2017)* |
| | | (A2) download date; version of data source | Published version dating from 2017 |
| | | (A3) basis of records | Forest inventories and research networks |
| | | (A4) spatial extent | Europe |
| | | (A5) temporal range | One period up to 2016 |
| | processing | (A6-1) duplicate coordinates | Dataset already cleaned |
| | | (A6-2) spatial/environmental outlier; error | Dataset already cleaned |
| | | (A6-3) spatial/coordinate uncertainty | All records on 1 km grid |
| | | (A7-1) sampling bias | Dataset arises from systematic sampling in forest inventories. Some countries are not taken up in the dataset. |
| | | (A7-2) spatial autocorrelation | Occurrence points were aggregated to the 1/24-degree spatial resolution of the climate data. |
| (B) Obtaining and processing environmental data | metadata & processing | (B1) source | TerraClimate (Abatzoglou *et al. 2018)* |
| | | (B2) download date; version of data source | Downloaded on 13/07/2022 |
| | | (B3) spatial resolution | 1/24-degree |
| | | (B4) temporal range | 1986-2015 |
| (C) Model calibration | data input | (C1) modeling domain | Climatic environmental space |
| | | (C2) number of background data | No background used, absences were derived from dataset. |
| | | (C3) sampling method for background data | / |
| | | (C4) variable selection | Parameters derived from three extreme variables that can limit species distribution. 16 other bioclimatic variables were included to adhere to common practice. |
| | algorithm | (C5) name | Generalised additive models (GAMs) |
| | | (C6) version of algorithm and software | gam function of the mgcv R-package version 1.8-42. |
| | | (C7) parameterization | The GAMS had a binomial distribution. The GAMs included a linear combination of smooth terms for all predictors. Smooth terms are penalised regression splines with 10 basis functions and smoothing parameter estimation using the generalised cross validation criterion. Model terms were not penalised. |
| | evaluation | (D1) evaluation index | Model AIC, AUC and edge AUC. |
| | | (D2) threshold for evaluation index | No threshold was used. |
| | | (D3) dataset used to evaluate models | AUC was based on a spatially blocked cross-validation design. |
| | output | (D4) format/transformation | Continuous probability of occurrence was used. |
| | | (D5) threshold | No threshold was used. |

| | | | |
|---|---|---|---|
| (D) Model transfer and evaluation | extrapolation | (D6) novelty of projected environments compared with training environments | Projectsion were only performed in areas inside the calibration area of the models. No projections to other time-periods were performed. |
| | | (D7) collinearity shift between training and projected environments | Not assessed. |
| | | (D8) extrapolation strategy | Projectsion were only performed in areas inside the calibration area of the models. No projections to other time-periods were performed. |
| | metadata | (D9) source | Not applicable |
| | | (D10) download date; version of data source | Not applicable |
| | | (D11) spatial resolution | Not applicable |
| | | (D12) temporal range | Not applicable |

#### Appendix I

#### Spatial predictions

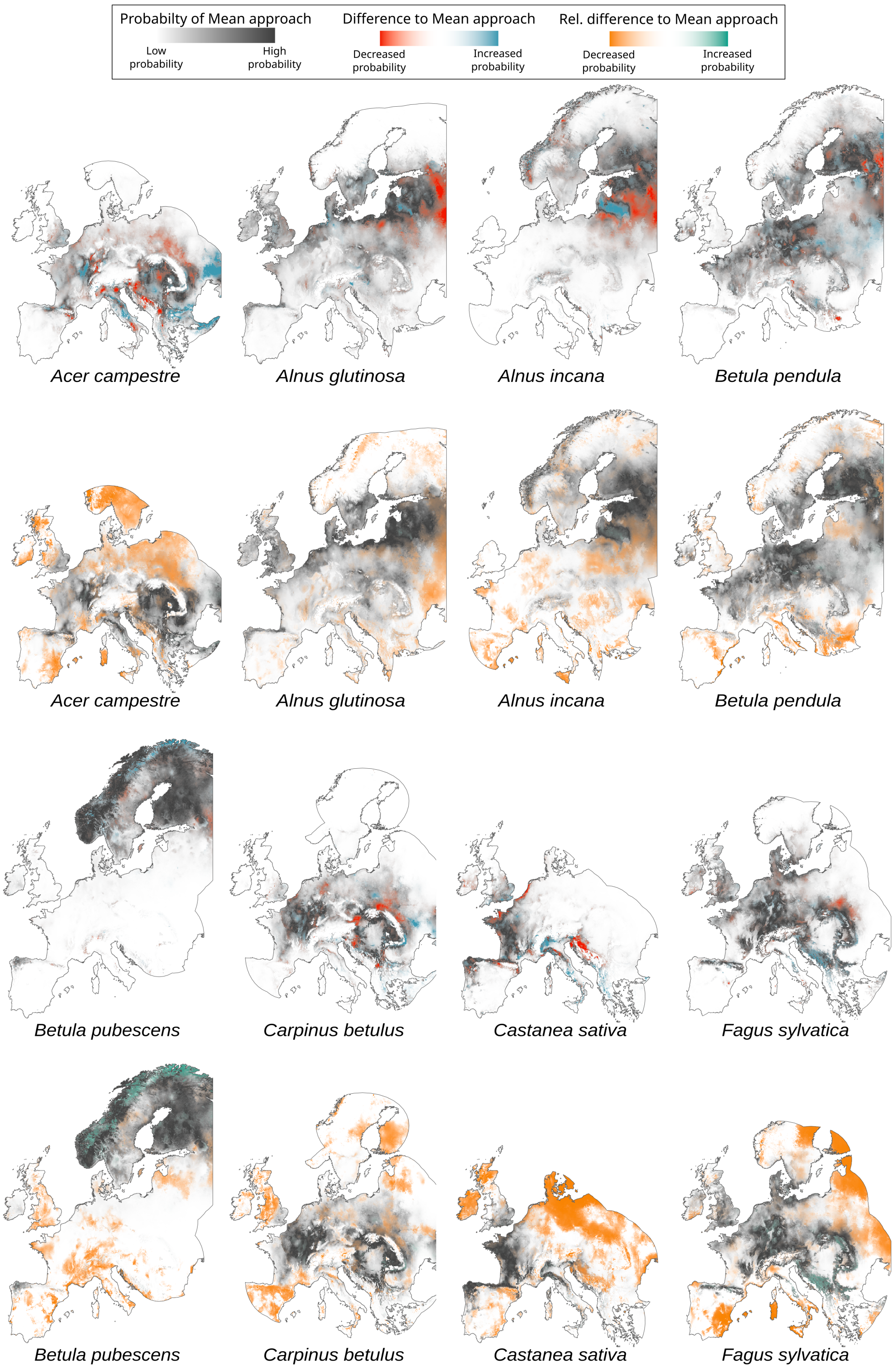


**Figure I1**

*A comparison between the spatial predictions derived from the GEV approach and the Mean approach are given for 8 tree species. The probability for the species as predicted by the Mean approach is given in a gradient from white to dark grey, while the difference in predicted probability between the GEV approach and the Mean approach is given with a gradient from red to blue. More opaque red indicates a greater decrease in probability while more opaque blue indicates a greater increase. Only the 500-km buffer around presences is shown. Predictions were performed on 1/24-degree spatial resolution.*

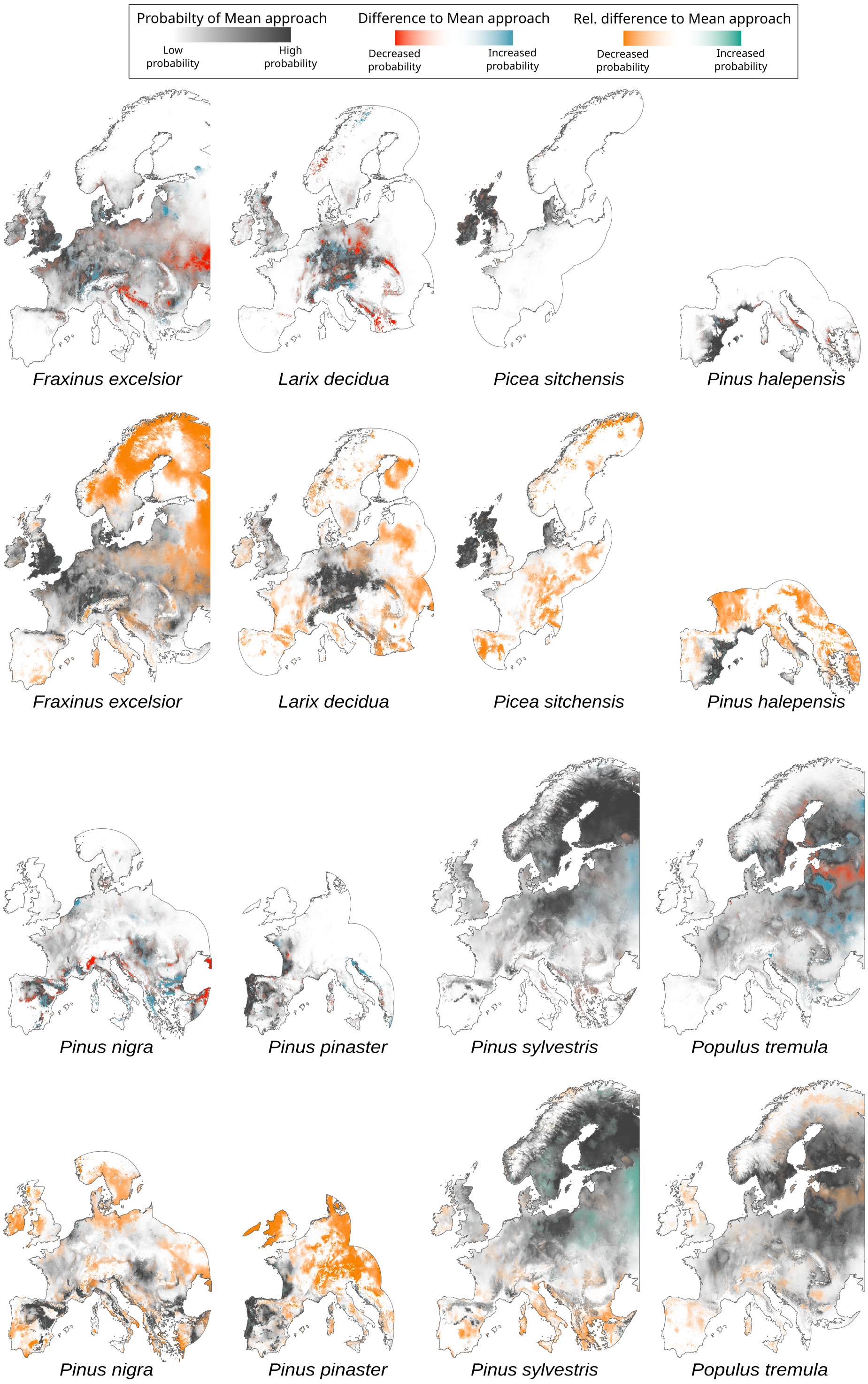


**Figure I2**

*A comparison between the spatial predictions derived from the GEV approach and the Mean approach are given for 8 tree species. The probability for the species as predicted by the Mean approach is given in a gradient from white to dark grey, while the difference in predicted probability between the GEV approach and the Mean approach is given with a gradient from red to blue. More opaque red indicates a greater decrease in probability while more opaque blue indicates a greater increase. Only the 500-km buffer around presences is shown. Predictions were performed on 1/24-degree spatial resolution.*

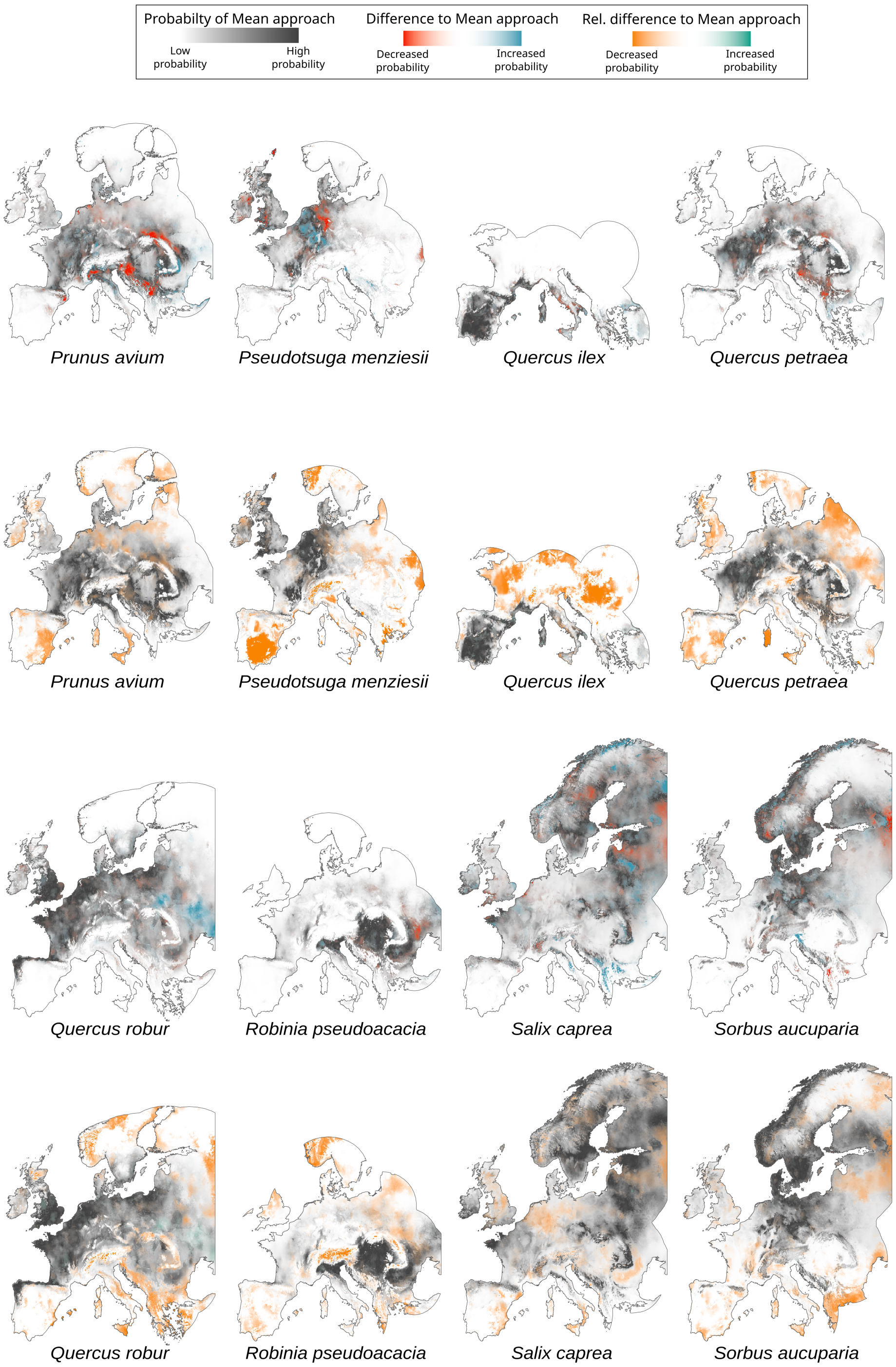


**Figure I3**

*A comparison between the spatial predictions derived from the GEV approach and the Mean approach are given for 8 tree species. The probability for the species as predicted by the Mean approach is given in a gradient from white to dark grey, while the difference in predicted probability between the GEV approach and the Mean approach is given with a gradient from red to blue. More opaque red indicates a greater decrease in probability while more opaque blue indicates a greater increase. Only the 500-km buffer around presences is shown. Predictions were performed on 1/24-degree spatial resolution.*
